# *PatientProfiler:* A network-based approach to personalized medicine

**DOI:** 10.1101/2025.01.31.635886

**Authors:** Veronica Lombardi, Lorenzo Di Rocco, Eleonora Meo, Veronica Venafra, Elena Di Nisio, Valerio Perticaroli, Mihail Lorentz Nicolaeasa, Chiara Cencioni, Francesco Spallotta, Rodolfo Negri, Francesca Sacco, Livia Perfetto

**Affiliations:** Department of Biology and Biotechnologies “Charles Darwin”, Sapienza University of Rome, 00185 Rome, Italy; Department of Statistical Sciences, Sapienza University of Rome, 00185 Rome, Italy; Department of Biology, University of Rome “Tor Vergata”, Rome, Italy; Ph.D. Program in Cellular and Molecular Biology, Department of Biology, University of Rome ‘Tor Vergata’, Rome, Italy; Institute of System Analysis and Informatics “Antonio Ruberti”, National Research Council (IASI-CNR), 00185 Rome, Italy; Istituto Pasteur Italia-Fondazione Cenci Bolognetti, Sapienza University, 00185 Rome, Italy; Institute of Molecular Biology and Pathology (IBPM), National Research Council (CNR) of Italy, 00185 Rome, Italy

**Author notes:** These authors contributed equally (Lorenzo Di Rocco, Eleonora Meo, Veronica Venafra).

## Abstract

Deciphering the intricate mechanisms underlying reprogramming in cancer cells is a crucial challenge in oncology as it holds the key to advance our ability to diagnose and treat cancer patients. For this reason, comprehensive and patient-specific multi-omic characterization of tumor specimens has become increasingly common in clinical practice. While these efforts have advanced our understanding of the molecular mechanisms underlying breast cancer progression, the identification of personalized therapeutic approaches remains a distant goal. The main shortcoming is the absence of a robust computational framework to integrate and interpret the available multi-dimensional data and to drive translational solutions.

To fill this gap, we developed *PatientProfiler*, a computational pipeline that leverages causal interaction data, annotated in our in-house manually-curated resource, SIGNOR, to address how the genetic and molecular background of single patients contributes to the establishment of a malignant phenotype. *PatientProfiler* is an open-source, R-based package composed of several functions that allows multi-omic data analysis and standardization, generation of patient-specific mechanistic models of signal transduction, and extraction of network-based prognostic biomarkers.

To benchmark the tool, we retrieved genomic, transcriptomic, (phospho)proteomic, and clinical data derived from 122 treatment-naïve breast cancer biopsies, available at the CPTAC portal. Thanks to this approach, we identified patient-specific mechanistic models (one patient, one network) that recapitulate dysregulated signaling pathways in breast cancer. This collection of models provides valuable insights into the underlying mechanisms of tumorigenesis and disease progression. Moreover, in-depth topological exploration of these networks has allowed us to define seven communities (subnetworks), each associated with a unique transcriptomic signature and a distinct prognostic value.

In summary, our work demonstrates that *PatientProfiler* is a tool for patient-specific network analysis, advancing personalized medicine towards the identification of actionable biomarkers and tailored therapeutic strategies.

## Introduction

The aim of personalized and precision medicine is to shape treatments over the molecular markers of a disease at single patient resolution^1^. Traditionally, precision oncology relies on histological, genomic, and transcriptomic biomarkers due to their scalability and cost-effectiveness. However, these approaches often result in generalized stratification and a subset of patients not benefiting from patient-drug matching^2^. Several observations suggest that cancer onset and progression cannot be solely explained by genetic alterations but involve a complex reprogramming of signaling pathways^3^.

Research showed that apoptotic pathway deregulation correlates with cancer cell survival advantage, leading to multidrug resistance^4^. Additionally, a study demonstrated that patients with similar pathway alterations, such as those in the PI3K/mTOR pathway, including high p4E-BP1 phosphorylation at specific sites (e.g., S65, T70), high pS6 (S235/236), and low pdcd4, are associated with worse recurrence-free survival (RFS) and overall survival (OS) clinical outcome in hormone receptor-positive breast cancer^5^.

These findings underscore the significance of data-driven models and pathway analysis in delivering more reliable and specific prognostic biomarkers, to guide therapeutic interventions in cancer research. As such, they support the need for a way to systematically address the signaling events dysregulated in individual patients and their employment to identify biomarkers.

As such, to achieve the milestone of personalized therapeutic solutions we need more specific biomarkers that consider the patient-specific signaling profile.

In recent years, to address the role of signal transduction in carcinogenesis, several independent groups^3^ as well as the Clinical Proteomic Tumor Analysis Consortium (CPTAC) invested in a comprehensive and multi-omic (e.g. genomic, transcriptomic, phosphoproteomic, etc.) profiling of cancer specimens^6^, offering the unique and unprecedented opportunity of unbiasedly and systematically assessing how cellular pathways are perturbed in a given malignancy, in general, and in individual patients.

Such advances in technology inspired the development of a range of bioinformatics tools capable of integrating these multi-omic layers to generate hypotheses about signaling mechanisms in the form of networks. The main hypothesis behind these approaches is that to understand carcinogenesis or cancer progression, we can use mechanistic/data-driven models. These models aim to describe experimental data via networks of regulatory interactions among proteins of the biological system under study. Consequently, they function as connectors to link genotypes to phenotypes at cellular, tissue, or organism levels.

To date, several computational methods combine untargeted omics data with prior knowledge to estimate the state of signaling networks in specific biological scenarios^7^. These methods vary in terms of input omics data, prior knowledge, and underlying methodologies.

Despite these efforts, no effective approach has yet been developed to identify the signaling mechanisms that are deregulated at the patient-resolution level, nor these have been effectively employed to nail down patient-tailored therapeutic regimens^8^.

In 2016, Drake et al contributed to the development of a pCHIPS^9^ by a pioneering approach, developing a tool that combines genomic, transcriptomic, and phosphoproteomic datasets from Prostate cancer patients to synthesize patient-specific networks highlighting druggable kinases. Generated networks, however, are undirected and do not inform about information flux propagation in the network, thus limiting the readability of the results.

More recently, Dugourd et al released COSMOS+, a tool that connects data-driven analysis of multi-omic data from publicly available cell line data with systematic integration of prior knowledge causal networks^10^. Recently we developed *SignalingProfiler*^11^, a computational strategy that combines genomic, phosphoproteomic, and transcriptomic data with networks of causal interactions derived from SIGNOR resource^12^ to extract personalized models of signal transduction.

Here we present *PatientProfiler*, a computational strategy that can: 1) handle and harmonize patient-specific data; 2) apply SignalingProfiler 2.0 to extract patient-specific protein activities and generate mechanistic networks linking them to cancer phenotypic hallmarks; 3) apply clustering studies to define stratification groups and 4) deliver novel and signaling-driven transcriptomic signatures.

As a use case, we applied *PatientProfiler* to deliver patient-specific mechanistic models of signal transduction for 122 breast cancer patients. We here show that by reducing data, complexity, and dimensionality *PatientProfiler* can provide valuable insights into the underlying mechanisms of tumorigenesis and disease progression. Moreover, in-depth topological exploration of these networks allowed us to define seven communities (subnetworks), each associated with a unique transcriptomic signature and a distinct prognostic value.

We here claim that *PatientProfiler*, by addressing sample specificity will pose the basis to bring proteogenomic studies into a translational environment.

## Methods

### Pipeline Description

We introduce *PatientProfiler*, an R-based workflow composed of two main parts: The first part aims to generate patient-specific models from multi-omic data (Steps 1-3). Whereas the second part leverages the so-generated models to deliver biomarkers of the disease (Step 4-5).

The code, the documentation and a tutorial are available at: https://github.com/SaccoPerfettoLab/PatientProfiler

#### Step 1, Dataset manipulation

Here, *PatientProfiler* harmonizes omic data through the “omics_update” function, which i) parses the data to make it compliant with further steps in the pipeline; ii) filters and handles missing data. As an example, genes/proteins exceeding a maximum percentage of missing data are removed; iii) performs quality checks on annotated sequences in phosphoproteomics and on identifiers and symbols used. Briefly, to ensure consistency, it updates the gene names and phosphosites according to the most updated UNIPROT sequence. It further adds the UNIPROT IDs and retrieves the peptides’ sequence window (i.e a 15-mer peptide centered on the phosphorylated amino acid); iv) performs sample-wise z-scoring to estimate the change in abundance of each analyte, in each sample. Importantly, applying the analyte-wise (by row) and/or the sample-wise (by column) z-score. The user can choose whether to use median- or mean-centered z-score. Values greater than and less than +/-1.96 (i.e., +/-2 standard deviations, p-value < 0.05) are considered to be significantly modulated.

More Specifically, for transcriptomic data it is possible to apply filtering and z-score computation. For proteomics and phosphoproteomics imputation, quality check and z-score computation. For phosphoproteomics users can retrieve the sequence window.

#### Step 2-3, Protein activity inference and generation mechanistic models

In the second and third steps of *PatientProfiler*, we developed two wrapper functions for the SignalingProfiler 2.0 R package^11^: (i) “extract_protein_activity” function, which infers patient-specific protein activities modulations from multi-omics data combining footprint-based methods and PhosphoScore algorithms, with default SignalingProfiler 2.0 parameters; (ii) “create_network” function, which builds patient-specific mechanistic models connecting each patient’s functionally annotated mutations to the inferred signaling proteins. This function internally calls five *ad hoc* developed functions to choose: a) PKN type, b) naive-network layout, c) CARNIVAL algorithm type, d) PhenoScore algorithm implementation, and e) formatting options for Cytoscape visualization. To simplify the network construction step (Step 3), we provide the “initialize_net_default_params” function, which automatically sets *default* parameters (PKN with direct interactions, two-layered naive networks, inverse CARNIVAL algorithm).

To generate cohort-wide protein activities and network collections, we implemented ‘extract_cohort_activity’ and ‘create_cohort_networks’ functions. These functions iteratively call the protein activity inference and network creation algorithms for each patient, resulting in a table of transcription factors, kinases/phosphatases, phosphorylated proteins with modulated activity, as well as a list of networks (1 network-1 patient) for the cohort.

#### Step 4: Network-based patient Stratification

This step leverages the collection of patient-specific mechanistic networks 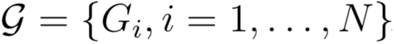 to simultaneously uncover patient clusters and extract the specific pathways highly associated with them. To this end, we design a pipeline consisting of four modules: 1) Networks Aggregation. In this module, we merge the collection of networks 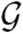 into a single aggregated network *G*\*. We focused exclusively on pathways from *G*\* originating from an activated node; 2) Bipartite Projection. The second module builds a bipartite graph *B* = (*V*, *E*) to model the patient-interaction relationship. In this graph, the vertex set is defined as 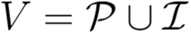, where 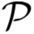 represents the set of patients in the cohort and 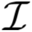 denotes the set of interactions occurring in *G*\*. An edge *e* = (*p*, *i*) exists between a patient node 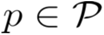 and an interaction node 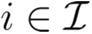 if and only if the corresponding patient *p* includes the interaction in their interactome. This topological structure enables us to link patients with common regulatory interactions. 3) *Community detection.* The third module involves detecting communities in *B*. To achieve this, we used the Louvain algorithm^13^, which applies greedy optimization techniques to iteratively refine clusters. This hierarchical approach begins by grouping nodes into communities that maximize modularity. These communities are then aggregated into super-nodes, and the process is repeated on the resulting graph. The algorithm continues this iterative optimization until no further improvements or refinements can be achieved; and 4) *Identification of Communities’ Networks.* In the last module, we identify the specific pathways associated with each patient community. This is achieved by assigning to each community the set of interactions of *G*\* grouped within the same cluster on the bipartite graph *B* in the previous step.

Additional details are available in Supplementary data.

#### Step 5, identification of biomarkers

In this last step, *PatientProfiler* offers the user the chance to derive transcriptomic signatures from the obtained communities. For each community detected in Step 4, *PatientProfiler* first retrieves all the patients belonging to the community and extracts their transcriptomic profiles. Next, for each gene performs an ANOVA followed by Tukey test and filters genes that are i) up-regulated in the patients belonging to the community (mean expression in the community > 0); ii) that display the highest variance in respect to the rest of the patients (background) (ANOVA adjusted p-value < 0.01); and iii) that have a minimum expression difference of 0.7 (mean expression in the community - mean expression in the background).

#### Network visualization

Networks were visualized in Cytoscape using the “RCy3” package (version 2.24.0) and the “createNetworkFromIgraph” function. The style “SP_pheno_layout.xml” was applied for the patient-specific mechanistic models.

### Breast Cancer use-case

#### CPTAC Data Collection (discovery cohort)

We retrieved Breast cancer data^14^ using the *cpatc* Python package (version 1.5.0rc1) with the “cptac.download” function and the parameter “dataset = Brca”. We downloaded transcriptomic, proteomic, phosphoproteomic, somatic mutation and clinical datasets. Transcriptomic, proteomic and phosphoproteomic data were analysed using Step1 of *PatientProfiler* (*Dataset manipulation)*, with default parameters. Briefly, we filtered data and computed the sample-wise (by column) z-score (median-centered).

Somatic mutation data was annotated as gain-of-function (GOF) or loss-of-function (LOF) using OncoKB API (https://www.oncokb.org/). In particular, we used MafAnnotator.py OncoKB tool via a command line with Python 3 to annotate the impact of each mutation on protein function (1 for GOF and −1 for LOF) (Supplementary Table 1).

To define patients’ subtypes we referred to the Non Negative Matrix Factorization clustering (NMF.Cluster) described in the publication by Krug et al., 2020, from which we extracted the data.

To generate the collection of mechanistic models and extract the communities, we exploited Step 2, Step 3 and Step 4 of *PatientProfiler.* Specifically, for the Step 2 we used the parameters hypergeom_corr = FALSE for the analysis = ‘tfea’ with correct_proteomics = FALSE, and for the analysis = ‘ksea’ with correct_proteomics = TRUE. For Step 3 and Step 4 we used default parameters.

To extract the transcriptomic signatures, we used patients’ communities (or subgroups) and patients’ transcriptomic profiles and employed Step5 of *PatientProfiler*, with default parameters for community. Only for communities 1, 5 and 6 (CL1, CL5 and CL6) we selected a minimum expression difference of 0.5 (mean expression in the community - mean expression in the background).

#### TCGA Data Collection (Validation cohort)

We downloaded transcriptomic and follow-up data for 1,094 breast cancer patients from the TCGA using the “TCGAbiolinks package” (version 2.32.0, November 2024). We first applied log(x+1) transformation and then we applied Step 1 of *PatientProfiler* (*Dataset manipulation)* to filter data and to compute the sample-wise (by column) z-score (median-centered). We only considered coding transcripts and rank them in descending order. Next, we used so-manipulated transcriptomic data to seek for the enrichment of each of the seven transcriptomic signatures (CL1-7). To this aim, we performed a gene set enrichment analysis using the “fgseaMultilevel” function with the parameter scoreType = ‘std’ and selected the signatures with lowest adjusted p-value. In this way, each patient from TCGA was assigned to a community. Finally, survival curves were generated using the “ggsurvfit” package (version 1.1.0).

### Over-representation analysis (ORA)

Over-representation analysis was performed using “gprofiler2” software (version 0.2.3) using the default background and selecting pathway terms from KEGG^15^, Reactome^16^ and Wikipathways^17^, biological processes from Gene Ontology^18^ and complexes from CORUM^19^. Bonferroni-adjusted p-values < 0.05 were considered significant. Only top enriched 10 terms were considered.

### Cell cultures

The human breast cancer cell lines MCF7, MDA-MB-361 and MDA-MB-231 were cultured at 37 °C in a 5% CO_2_ atmosphere. Mycoplasma contamination was assessed monthly (Abm, US). MCF7 and MDA-MB-231 cells were grown in high-glucose DMEM with sodium pyruvate (Corning, US), containing 10% FBS (Corning, US) and 1 mmol/L l-glutamine (Sigma-Aldrich, DE). MDA-MB-361 cells were grown in RPMI 1640 (Corning, US), containing 10% FBS (Corning, US) and 1 mmol/L l-glutamine (Sigma-Aldrich, DE).

### Protein purification and quantification

Cell lysis was carried out in RIPA Buffer (ThermoFisher Scientific, US) in the presence of protease inhibitor cocktail (Complete, Roche, US), phosphatase inhibitors (Cocktail Tablets, Roche, US) according to the manufacturer’s protocol guidelines. An additional phosphates inhibitor β-Glycerophosphate (10mM) (Sigma-Aldrich, DE) was also added to the lysis buffer. Samples have been purified with centrifugation @16000g 4°C for 30 minutes. Proteins were quantified using the Bradford Protein Assay Reagent (ThermoFisher Scientific, US).

### Western blot

For each sample 20 µg of total protein lysate has been loaded on a denaturing 10% PAGE of acrylamide/bisacrylamide solution 29:1 (Serva, DE). Page Ruler Protein Ladder (ThermoFisher Scientific, US) was used. Proteins were transferred to 0.4µm nitrocellulose membranes (Bio-Rad, US) at 90V for 90 minutes. The transfer efficiency was verified using the Ponceau S solution (PanReac, ES, USA). Membranes were blocked in TBS containing 0.1% Tween-20 (T-TBS) (Bio-Rad, US), 5% milk (Sigma-Aldrich, DE), or 5% (bovine serum albumin BSA (Sigma-Aldrich, DE). The primary Antibodies used are: anti-CDK5 (Cell signaling, 2506) 1:1000, anti-Phospho-CDK5 (Tyr15) (Thermo, PA577909) 1:1000, anti-Phospho-CDK2 (Thr160) (Cell signaling, 2561) 1:1000, anti-vinculin (ab129002, Abcam) 1:1000 in 5% BSA-TBST; anti-CDK2 (SantaCruz, sc-6248) 1:500 and anti-Cyclin A (B-8) (SantaCruz, sc-271682) 1:500 in 5% skim milk-TBST. Goat anti-rabbit HRP conjugate (1:10000, Thermo Fisher Scientific) and Goat anti-mouse (1:10000, Thermo Fisher Scientific) were used as secondary antibodies. Chemoluminescence was acquired using Clarity western ECL substrate (Bio-Rad, US) in a ChemiDoc MP Imaging System (Bio-Rad, US). For images quantification ImageLab Software has been used.

## Results

### *PatientProfiler*, Pipeline Overview

*PatientProfiler* is a novel R workflow designed to unbiasedly integrate cancer-derived multi-omic data at the patient-resolution level. The scope of *PatientProfiler* is to get mechanistic insight into pathways and molecular mechanisms that are deregulated in individual patients, giving as output network-based prognostic biomarkers, and, possibly, formulation of novel therapeutic strategies in oncology.

The entire pipeline is freely accessible and available for reuse and interoperability at https://github.com/SaccoPerfettoLab/PatientProfiler.

Here we provide a step-by-step description of the method, which consists of five key steps, organized into two main parts. The first part aims to generate patient-specific models from multi-omic data (**Figure 1, Steps 1-3**). Whereas the second part leverages the so-generated models to deliver biomarkers of the disease (**Figure 1, Step 4-5**).

**Figure 1.**
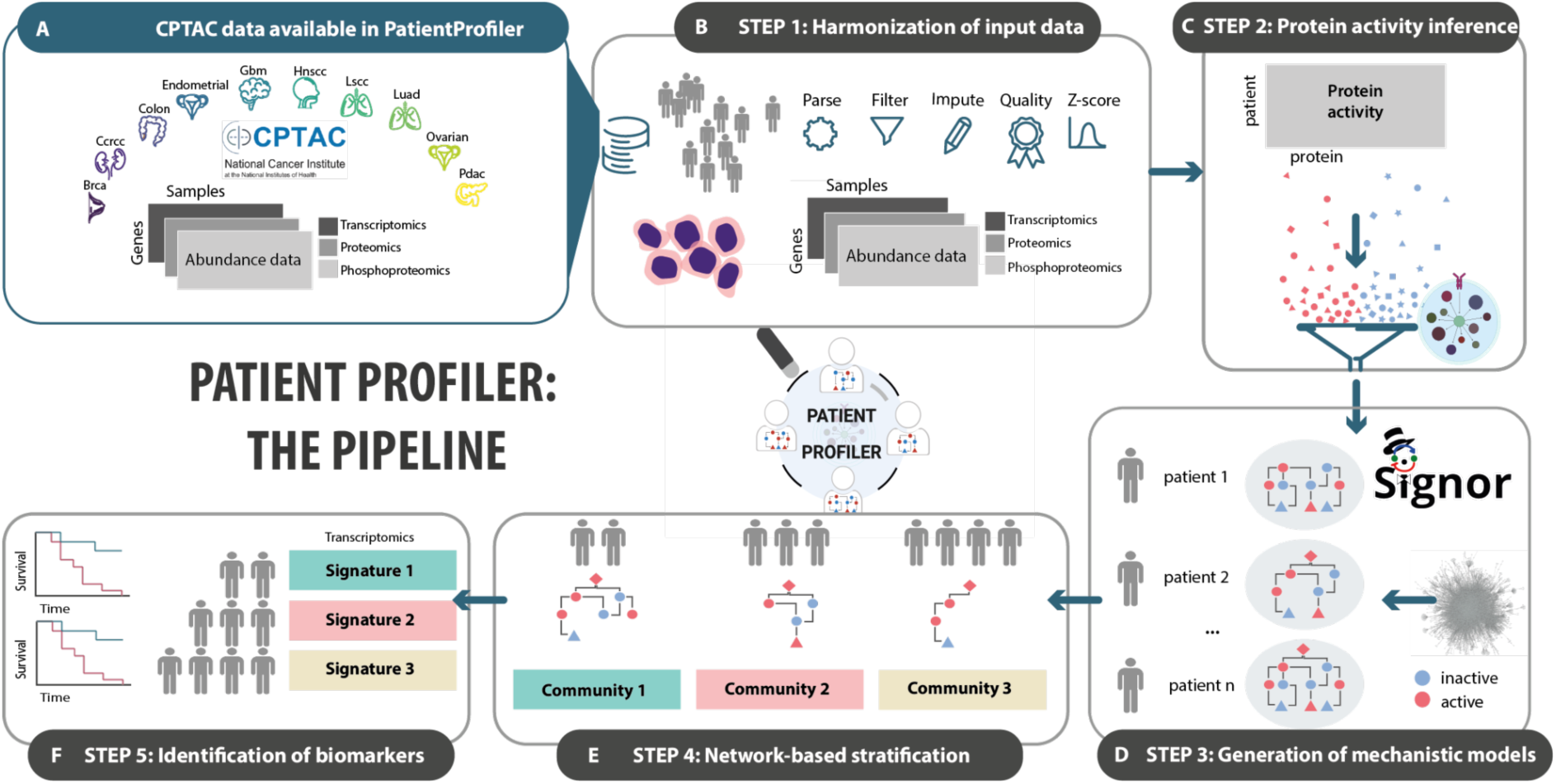
*PatientProfiler*, the pipeline. **A)** Summary of CPTAC data^6^ available in *PatientProfiler*. **B)** STEP1 allows users to parse custom data or data imported from CPTAC. Data accepted: transcriptomics, proteomics, and phosphoproteomics from multiple samples. **C)** In STEP2 PatientProfiler allows the user to unbiasedly estimate, within each sample, protein activities from the multi-omic data (red: active proteins; blue: inactive proteins). **D)** In STEP3 *PatientProfiler* leverages causal data in public resources^12,20^ to hierarchically connect protein activities from STEP 3 and to build for each sample (patient) a mechanistic model. **E)** In STEP4 *PatientProfiler* leverages mechanistic models from STEP3 to detect communities based on network structural similarities. Each community is represented by a group of samples (patients) and by a subgraph capturing key signaling interactions within the group. **F)** In STEP 5, communities detected in STEP4 are used in combination with relative gene expression profiles to identify biomarker transcriptomic signatures.

#### Step 1, harmonization of input data

The main bottlenecks toward the application of bioinformatics tools derives from the heterogeneity of available data, which often is sparse, outdated, inconsistent, and distributed in different formats. The first module of *PatientProfiler* offers optional functions that ensure and facilitate standardization of input transcriptomic, and (phospho)proteomic data. While this step was optimized to access cancer data available at the CPTAC portal, we generalized it to allow users to analyze custom information from cancer cell lines or tissue biopsies.

In summary, *PatientProfiler* can take as input multi-dimensional data (multiple omic levels, multiple patients, multiple analytes), yet analyzing each patient individually. This feature ensures that batch effect is not relevant since patient data are analyzed internally and future patients can be analyzed individually. To note, users can select as input in *PatientProfiler* a user-compiled MOFA analysis as well as choose among a number of pre-processed input data. At the time of writing *PatientProfiler* grants access to pre-harmonized proteogenomic data from more than 1600 patient-derived samples, organized in 10 different tumor types, derived from CPTAC data, for access and analysis of future users (**Figure 1A**).

#### Step 2, protein activity inference

As a second step, *PatientProfiler* unbiasedly derives the activity of proteins from the integration of pre-processed data with prior knowledge information deposited in public repositories. More in detail, *PatientProfiler* leverages the footprint-based and the PhosphoScore approaches implemented in our recently developed SignalingProfiler 2.0^11^, to estimate the activity of kinases, phosphatases and phosphorylated proteins from the phosphoproteomic dataset; and the activity of transcription factors from the transcriptomic dataset.

The result of this step is a list of patient-specific signaling proteins for which we can observe a change in activity in the diseased tissue.

#### Step 3, generation of mechanistic models

As a further step, *PatientProfiler* leverages SignalingProfiler 2.0 i) to retrieve from our SIGNOR repository causal interactions that can explain the experimentally observed changes in protein activities; and ii) to estimate the activation level of hallmark phenotypes (e.g. Apoptosis, Proliferation). These interactions are then hierarchically organized to link deregulated proteins, derived from Step 2, to resulting phenotypes. The outcome of this step is a collection of coherent networks portraying the molecular mechanisms underlying patient-specific disease development.

So generated networks can be directly visualized in Cytoscape, in two formats: the first displays the complete generated network; ii) the second highlights the circuits that lead to the specific deregulation of user-selected phenotypes.

Importantly, thanks to the usage of visual features (e.g., color and style of the nodes and edges), these networks facilitate the interpretation, while reducing complexity and noise, of the input data.

#### Step 4, network-based stratification

The essence of the second part of *PatientProfiler* (**Figure 1, Step 4-5**) is to identify network-based features that can explain differences among groups of patients, thus providing prognostic biomarkers. In this perspective, patients are usually stratified by cancer type (e.g. in Pan-Cancer analyses), by histologically-defined subtypes, or by other clinical features and metadata^21^. These stratifications clearly depend on the availability in the source dataset. The premise behind *PatientProfiler* is that patients displaying similar deregulated pathways are likely to display similar clinical outcomes and should be treated similarly. *PatientProfiler* offers the opportunity to run agnostic clustering analysis, to identify novel stratification groups (communities) associated with common network modules.

In this step, *PatientProfiler* first builds a bipartite graph incorporating every patient-interaction relationship. This bipartite structure enables the analysis of patient connectivity through shared regulatory features. Dense clusters of nodes within the graph reveal groups of patients that are highly interconnected through specific subsets of interactions. For this reason, we employed the Louvain algorithm, which applies greedy optimization techniques to iteratively refine clusters. This hierarchical approach begins by grouping nodes into communities that maximize modularity. Importantly each resulting community is represented by i) patient members sharing common signaling modules; and ii) subnetwork of interactions significantly connecting these patients within the bipartite graph.

#### Step 5, identification of biomarkers

In this last step, *PatientProfiler* offers the user the chance to derive transcriptomic signatures to use as biomarkers. The idea is that the communities obtained in Step 4 allow us to stratify patients based on a similar rewiring of the signaling. Here, *PatientProfiler* exploits an ANOVA + Tukey test to identify transcripts whose overexpression is associated with the community. The identification of biomarker transcripts is particularly important since transcriptomic datasets are more common in clinical practice, where at the current moment only a very limited number of patients can be profiled at the proteogenomic level. We believe that these network-driven transcriptomic signatures properly reflect the underlying signaling rewiring, which is driving each subgroup unique clinical behavior and, as such, can be used to stratify and classify novel patients.

### Breast Cancer use case

As a use case, we set out to analyze with *PatientProfiler* proteogenomic data derived from a cohort of 122 breast cancer patients^14^, available at the CPTAC portal. The scope is to demonstrate that the tool can provide insights into the patient-specific cell reprogramming that underlies breast cancer and use this information to stratify patients and deliver novel biomarkers (**Figure 2A**).

**Figure 2.**
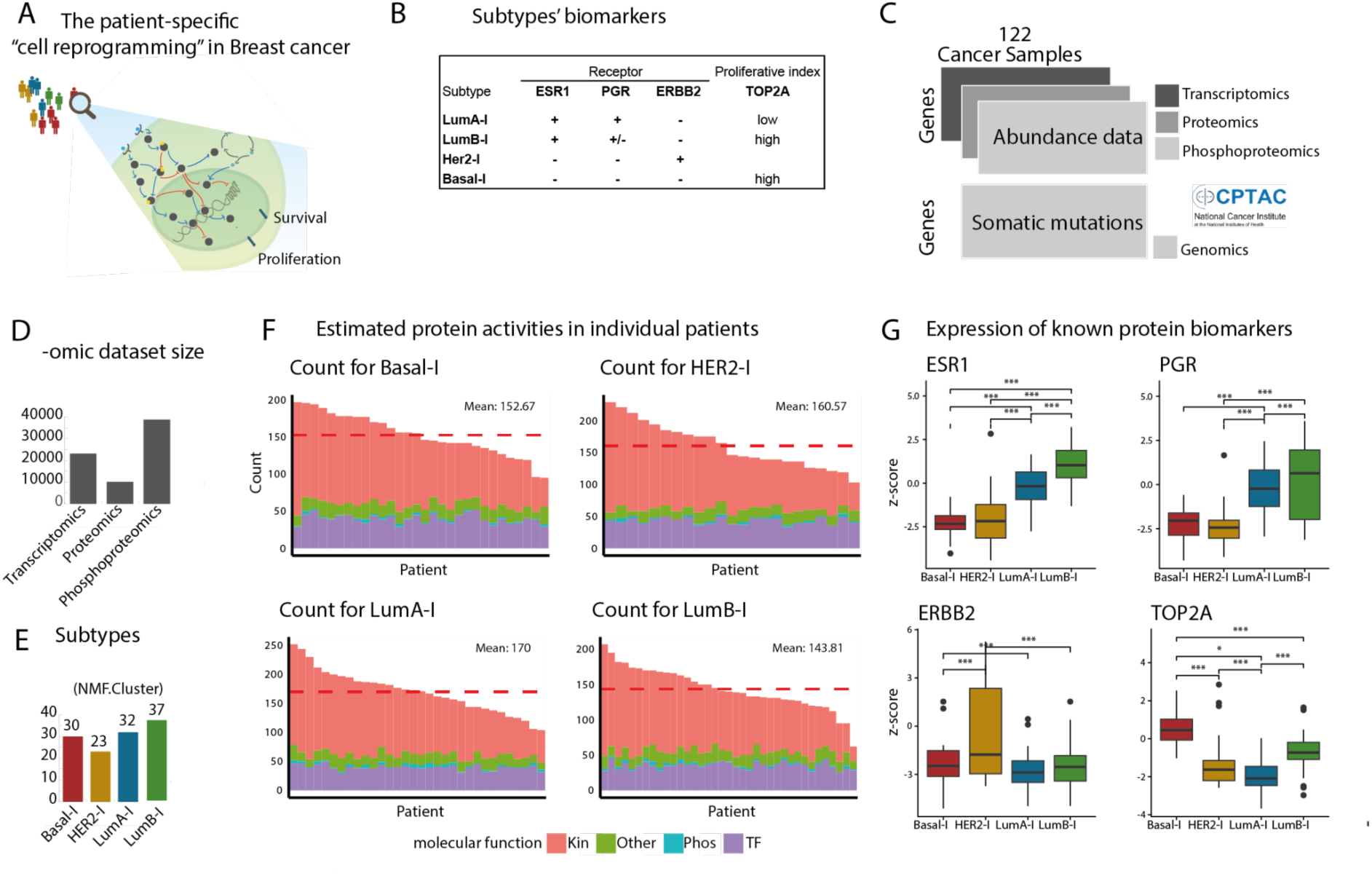
Breast cancer use case, STEP 1-2. **A)** Cartoon representing the scope of the use case. **B)** Breast cancer subtypes: Traditional subtyping include Luminal A-like tumors (LumA-l)(ESR1+, PGR+ and ERBB2-, low proliferative index); Luminal B-like tumors (LumB-l)(ESR1+, PGR+/- and ERBB2-, high proliferative index); HER2-like tumors (HER2-l)(ESR1-, PR- and ERBB2+); and Triple-negative breast cancer or Basal-like (Basal-l)(ESR1-, PGR- and ERBB2-, high proliferative index) whose features include high invasiveness, high metastatic potential, proneness to relapse, and poor prognosis. **C)** Breast cancer data was retrieved from *PatientProfiler* or from the CPTAC portal using the jupyter ‘cptac’ package^31^. **D)** Omic level size, relative to transcriptomic, proteomic, and phosphoproteomic data from a cohort of 122 breast cancer patients^14^ (**C**). **E)** Classification in subtypes (Basal-l, LumA-l, LumB-l, and HER2-l) and relative group size, as per original publication and referring to the NMF.Cluster^14^ **F)** Estimated protein activities in individual patients, stratified by subtypes. The dashed-red bar indicates the average amount of estimated proteins in each subgroup. **G)** Protein expression level of known biomarkers in different subtypes, after data manipulation (STEP1) (ESR1: estrogen receptor, PGR: progesterone receptor, ERBB2/HER2: human epidermal growth factor receptor 2, TOP2A: proliferative biomarker). Red, blue, purple, and green bars refer to kinases, phosphatases, transcription factors, and other types of proteins, respectively. Subtype color code: Basal-l in red, LumA-l in blue, LumB-l in green, and HER2-l in yellow.

Very briefly, Breast cancer is one of the most commonly diagnosed cancers among women and the second leading cause of cancer death among women overall^22^. Breast cancer is a highly heterogeneous disease under genetic, molecular, histopathological, and clinical terms. For prognostic prediction and treatment decision-making, breast tumors are traditionally classified into subtypes, as defined by immunohistochemistry or mRNA levels of estrogen receptor (ESR1), progesterone receptor (PGR), human epidermal growth factor receptor 2 (ERBB2/HER2), and proliferative biomarkers Ki67and TOP2A^23^(**Figure 2B**).

Traditional subtyping includes Luminal A-like tumors (LumA-l), Luminal B-like tumors (LumB-l), HER2-like tumors (HER2-l), and Basal-like (Basal-l), that is a Triple-negative breast cancer with high invasiveness, high metastatic potential, proneness to relapse, and poor prognosis. Because Basal-l tumors lack ESR1, PGR, and HER2 expression, they are not sensitive to endocrine therapy or HER2 treatment, and standardized Basal-l treatment regimens are still lacking^24^.

The heterogeneity of breast cancer biology deeply challenges the drive for personalized treatment and the present subtype classification remains insufficient in many cases^25^. For instance, within the LumB-l and Basal-l groups, we can observe substantial molecular diversity that influences therapeutic response and clinical outcomes^26^. Recent studies highlighted the need of further stratification, especially within these subtypes to identify clinically relevant molecular features and guide tailored treatment strategies^27,28,29,30^.

To this scope, we accessed CPTAC data available in *PatientProfiler* (March 2023) and extracted transcriptomic, proteomic, and phosphoproteomic datasets, displaying analyte abundance of 23,121 genes, 10,107 proteins, and 38,775 phosphosites respectively, in 122 breast cancer patients (**Figure 2C-D**). In addition, we also retrieved somatic mutation and clinical information^14^. As displayed in **Figure 2E**, patients in the cohort are homogeneously distributed across the four breast cancer subtypes: Basal-l, HER2-l, LumA-l, and LumB-l.

Most frequent mutation events in the cohort appear to be loss-of-function (LOF) and gain-of-function (GOF) mutations in the TP53 and PIK3CA genes, respectively (**Supplementary Figure 1, Table 1**). Interestingly, while the mutation in TP53 seems to associate prevalently with LumA-l (Chi-square p-value: 0.001) and Basal-l patients (Chi-square p-value: 1.02 e-07), the combination of the two is enriched in LumA-l patients, exclusively and tends to anti-correlate with Basal-l patients (Chi-square p-value: 0.003). However, the remaining mutations have only a mild effect on patient stratification. Next, we applied data-preprocessing functions, implemented in *PatientProfiler* to harmonize the data, and, for each patient and each dataset, we computed the sample-wise z-score as a proxy of entity (phosphosite, transcript, etc.) change in abundance.

Importantly, data manipulation and normalization preserve the differences between subgroups, especially at the proteome and transcriptome levels (**Supplementary Figure 2**). Also, the expression of known subgroup-associated biomarkers (ESR1, PGR, ERBB2, and TOP2A) is in line with the classification, both at the protein and transcript levels (**Figure 2B and G, and Supplementary Figure 3A**). In fact, ESR1 and PGR proteins appear to be significantly upregulated in LumA-l and LumA-l patients, with greater variability in the latter, ERBB2 in HER2-l, whereas Basal-l patients display lower levels of the three receptors. In line, Basal-l and LumB-l show higher levels of the proliferative TOP2A marker.

In summary, the tool correctly maintains differences between classical subtypes and has the potential to reflect their distinct molecular profiles.

### Application of *PatientProfiler* to Breast cancer data

#### Protein activity estimation

Analyte abundance is usually used to identify biomarkers, however, this type of data tends to be noisy and too complex to interpret^32^. To reduce dimensionality and complexity, we exploited the second step of *PatientProfiler* (**Figure 1C**), which leverages footprint-based and PhosphoScore analyses in SignalingProfiler 2.0, to infer the activity of proteins from multi-omic information (**Figure 2F-G**). In particular, for each patient in the cohort, we combined prior-knowledge information, namely kinase- and phosphatase-phosphosite pairs (also known as ‘regulons’) extracted from resources such as SIGNOR and PhosphoSitePlus and from large in vitro experiments^33^, with the relative change in abundance of the target phosphosites, as derived from the phosphoproteomics, to estimate the activity of kinases and phosphatases (**Figure 2F**, Kin and Phos), respectively. Similarly, we used transcription factor-target gene pairs from SIGNOR and Dorothea in combination with the transcriptomics to estimate the activity of transcription factors (**Figure 2F**, TF). Finally, by exploiting the PhosphoScore function implemented in SignalingProfiler 2.0, we exploited the regulatory role of specific phosphosites on the target protein to infer changes in activity. Importantly, the PhosphoScore function permits to extend the prediction also to other types of proteins (**Figure 2F**, Other).

Thanks to this approach, we were able to predict the activity of 712 proteins in the entire cohort (**Supplementary Figure 4, Supplementary Table 2**).

Importantly, protein activity inference preserves the nature of the subtypes where the activity profile of known subgroup-associated biomarkers (ESR1, ERBB2, and TOP2A) is in line with expectations (**Supplementary Figure 3B**).

As shown, the number of predicted proteins appears quite homogeneous in the different subtypes (**Figure 2F**), with LumA-l patients displaying an average increased number (170) of predicted proteins. Of note, in the four groups, the majority of inferred proteins are kinases, followed by transcription factors; this might be due to a combination of factors which include from one hand the important role of these proteins in mediating signal transduction, and, from the other hand, a regulon size bias in the prior knowledge used.

Finally, we sought for proteins displaying the strongest difference in activities in different subgroups. To this aim, we performed one-way ANOVA and plotted top significant results. As shown in **Figure 3**, cell cycle-associated proteins, such as the cyclin-dependent kinases (CDKs) and the E2F4 oncogenes appeared to be strongly dysregulated between subgroups, displaying an increased activity in the most severe breast cancer subtype, Basal-l.

**Figure 3.**
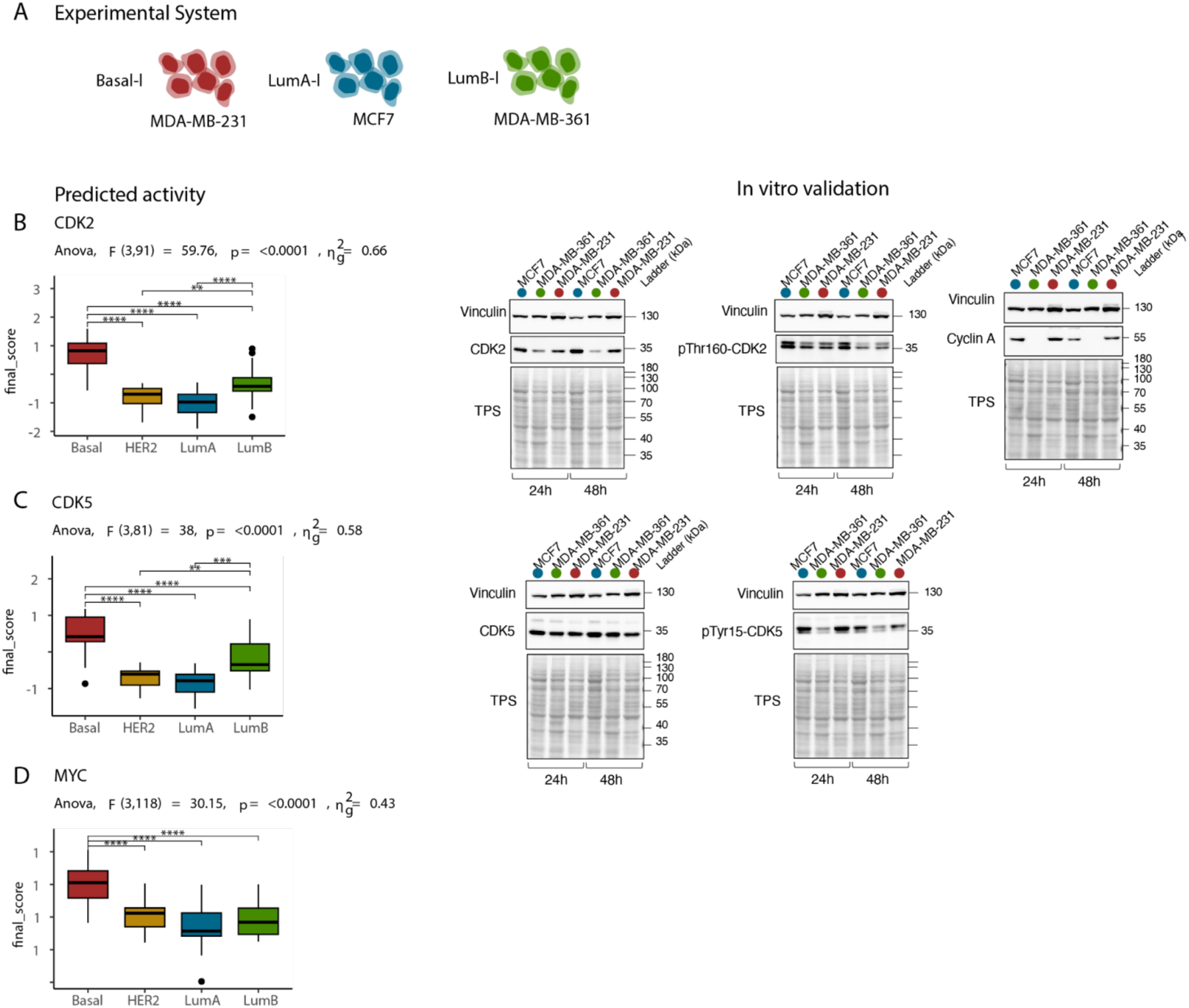
In vitro validation, STEP 1-2. **A)** Cartoon representing the experimental system: Basal-l, LumA-l and LumB-l subtypes were represented by MDA-MB-231, MDA-MB-361 and MCF7 cell lines, respectively. **B)** CDK2 activity (final_score) in different subtypes predicted by *PatientProfiler* (STEP2)(left panel) and relative experimental validation (right panel). Right panel: protein levels of CDK2, phosphorylation levels of activatory p-Thr160 CDK2 and protein levels of CyclinA in MDA-MB-231(red dots), MDA-MB-361 (green dots) and MCF7 (blue dots) cell lines after 24h and 48h were detected by western blot analysis. **C)** CDK5 activity (final_score) in different subtypes predicted by *PatientProfiler* (STEP2)(left panel) and relative experimental validation (right panel). Right panel: protein levels of CDK5, phosphorylation levels of activatory p-Tyr15 CDK5. in MDA-MB-231(red dots), MDA-MB-361 (green dots) and MCF7 (blue dots) cell lines after 24h and 48h were detected by western blot analysis. All the western-blotting results shown here were representative of three independent experiments. **D)** MYC activity (final_score) in different subtypes predicted by *PatientProfiler* (STEP2).

In line with these results, we observe that CDK2 and CDK5 are more active in Basal-l cell lines (MDA-MB-231) as compared to cell lines representative of other subtypes (Figure 3A-C).

Similarly, MYC and MYCN oncogene transcription factors display similar behavior, suggesting stronger levels of proliferation in this subtype, in line with previous evidence^34,35^.

In summary, inferred protein activities successfully reduced the complexity of the initial data, by increasing the level of abstraction and correctly representing breast cancer subtypes.

#### Network generation

A key challenge in multi-omics data integration is extracting the cause-effect relationships underlying the experimental data. Translated to our use case, this task attempts to address specific molecular events deregulated in individual patients, in the form of a signed and directed graph (**Figure 4A, Supplementary Figure 5**).

**Figure 4.**
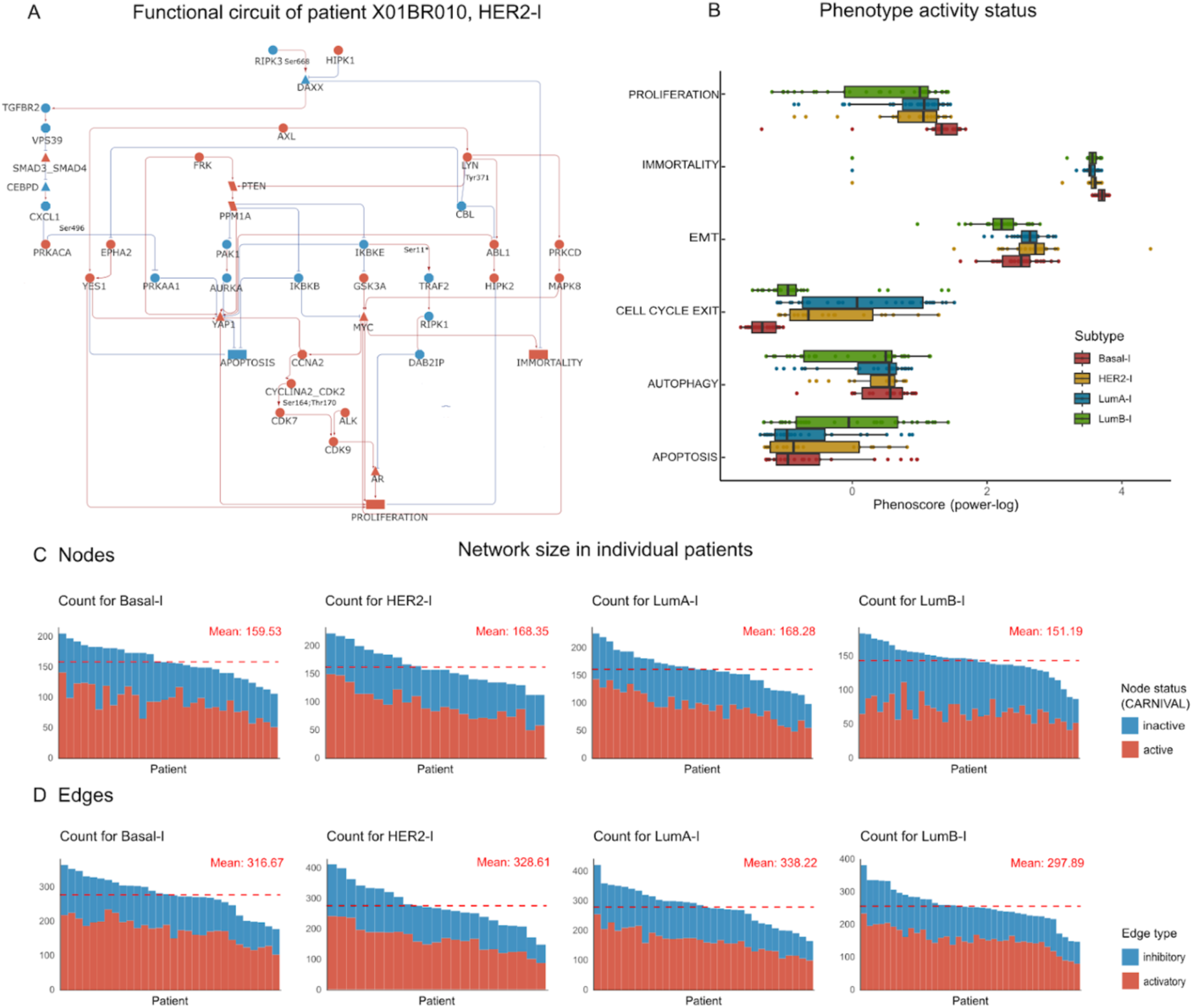
Breast cancer use case, mechanistic models generation (STEP 3). **A)** Example of a mechanistic model generated with *PatientProfiler*. The graph depicts the functional circuits to key phenotypes (rectangles) in the HER2-l patient X01BR010. Red and blue nodes represent active and inactive proteins, respectively; red arrows and t-shaped edges represent activatory and inhibitory relationships, respectively. Detected phosphorylated residues are shown as edge labels (e.g., Ser668). **B)** Activation level (power-log of the PhenoScore) of key hallmark phenotypes in different subtypes as measured in the STEP3 of the *PatientProfiler* pipeline. Subtype color code: Basal-l in red, LumA-l in blue, LumB-l in green, and HER2-l in yellow. **C-D)** Summary of the network size, as shown by the distribution of the number of nodes (C) and edges (D) in individual patients, stratified by subtypes. The dashed-red bar indicates the average amount of estimated nodes and edges in each subgroup.

Importantly, in each resulting network, nodes represent biological entities associated with an activity status (**Figure 4B-C, Supplementary Figure 6**) and edges the regulatory interactions, at the PTM-resolution level, among them (**Figure 4D**).

The result of this approach is a collection of 122 mechanistic networks built by unbiasedly integrating patient-specific molecular profiles with prior knowledge information. This collection can be accessed for analysis, visualization, and reuse via NDEx at the link https://www.ndexbio.org/#/networkset/bb975f9c-dcac-11ef-8e41-005056ae3c32?accesskey=5210ba80885b9ca10526b2fa52bfcf6643bb50e88401219c574796207db01b69

Importantly, for each patient in the cohort, we made available the full interactome generated and a subnetwork displaying the functional circuits impacting key cancer hallmarks (**Figure 4A**).

As displayed in **Figure 4A**, the so-generated networks permit the easy inspection of patient-specific circuits linking mutated proteins to dysregulated proteins and eventually to altered cancer hallmarks. Also, they allow comparison of patients and/or subtypes under different aspects such as network characteristics, and functional and topological properties.

These models are heterogeneous in size, as displayed by the number of nodes and edges which span from 100-200 and 150-400, respectively. To note, this variability is independent of the subtype. Also, the distribution of active/inactive nodes and activatory/inhibitory edges appears more homogeneous (**Figure 4C-D**). Interestingly, only few nodes in the network collection display similar activity in all patients (**Supplementary Figure 4 and 6**), whereas most entities display a heterogeneous activation pattern.

Similarly, hierarchical clustering indicates that although small groups of patients tend to cluster together, a more complex distribution of patients can be observed thus confirming that a certain level of heterogeneity characterizes patients within the same subgroup/subtype (**Supplementary Figure 4 and 6**).

Next, to determine whether the so-generated models could recapitulate the malignant behavior of the tumor samples as well as the different severity of the four subtypes, we set out to compare the activation levels of a number of cancer phenotypes^36^. As shown in **Figure 4B**, we could observe the tendency of model entirety to display increased levels of pro-oncogenic and pro-invasive phenotypes, namely the up-regulation of proliferation and epithelial-mesenchymal transition, along with the suppression of apoptosis and cell-cycle exit, indicating that models, in general, have the capability to recapitulate a malignant context. Also, in line with the expectations, Basal-l models (red bars in **Figure 4B**) are associated with more aggressive behavior for all considered phenotypes.

In summary, *PatientProfiler* provided for the first time a framework portraying biologically meaningful molecular mechanisms underlying patient-specific disease development. Networks facilitate the readability and interpretability of multi-omic data while reducing complexity and noise and offer novel features that can be used to determine similarities and differences between and within breast cancer subtypes. Overall, the network inspection suggests that part of the Basal-l models displays dysregulated cell-cycle progression, which is characteristic of a more aggressive cancer type^37^.

#### Communities’ prioritization and transcriptome signatures extraction

One of personalized medicine goals is the identification of more granular and specific biomarkers capable of stratifying patients at diagnosis in a more specific way, to better predict prognosis and therapeutic interventions ^38,39^. This is especially true in the case of breast cancer, where at diagnosis patients are assigned one subtype based on IHC and/or expression levels, with limited success in predicting disease outcome and drug response.

Since patients displaying similar deregulated pathways are likely to show a similar evolution of the disease as well as a similar response to therapies^40^, we established to use a community detection analysis to identify novel breast cancer subgroups, that might reveal novel network-derived prognostic biomarkers.

Leveraging the previously generated collection of regulatory networks, each capturing the molecular mechanism underlying the pathogenesis of a patient, we stratified the input cohort into distinct communities. Specifically, we employed the community detection approach (STEP 4 of *PatientProfiler*) to stratify the 122 patients of the cohort according to the topological structure of the generated models (**Figure 5A**).

**Figure 5.**
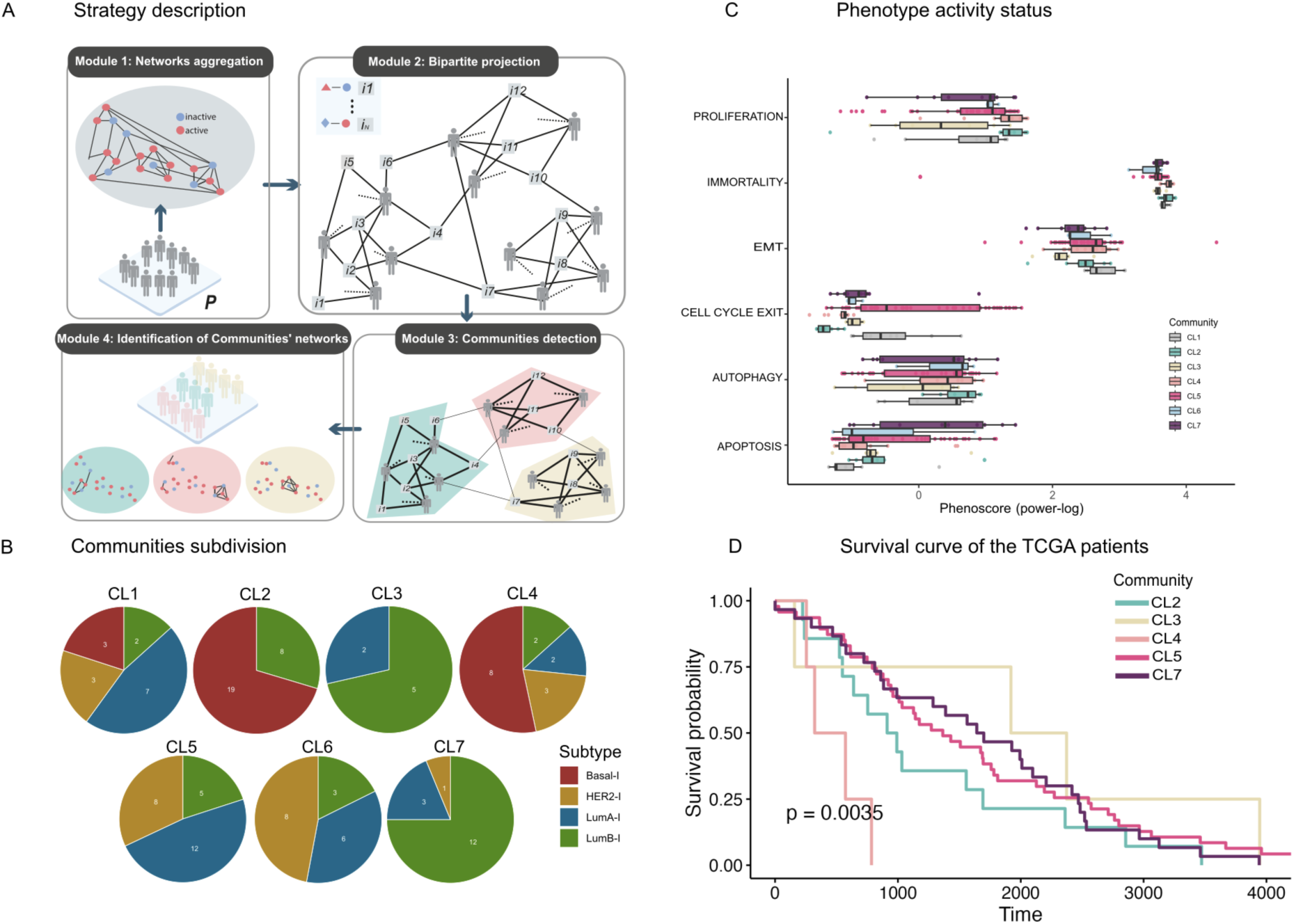
Community detection and their clinical impact (STEP 4). **A)** Cartoon depicting the community detection algorithm implemented in *PatientProfiler*, STEP4. Interactions from input patients are aggregated into a single network and projected into a bipartite patient-interaction relationship. Finally, modularity optimization is applied to identify distinct patient communities and their corresponding sub-interactomes. **B)** Our cohort of 122 Breast Cancer patients is divided into seven communities (CL1-7). Pie charts depicting the subtype composition of each community. Subtype color code refers to the NMF. Cluster classification, as derived from the original publication^14^: Basal-l in red, LumA-l in blue, LumB-l in green, and HER2-l in yellow. **C)** Activation level (power-log of the PhenoScore) of key hallmark phenotypes in different communities as derived from the mechanistic models. Community color code is summarized in the legend. **D)** Kaplan-Meier plot showing survival probability of Breast Cancer patients from The Cancer Genome Atlas (TCGA)^41^ stratified in communities.

By this approach, we were able to identify seven communities of patients (**Figure 5B, Table 3,** NDEx link: https://www.ndexbio.org/#/networkset/b03ab6d1-dcaf-11ef-8e41-005056ae3c32?accesskey=d974e5ac1f418cca58857dcf28044b9f9122c2c0e058980872e47c34f8cc380f) sharing common network topological structures (**Figure S10 – community networks**). These communities appear heterogeneous and include different subtypes, even if each of them is characterized by the overrepresentation of one subtype over the others. For example, Community 2 and Community 7 display a significant overrepresentation of Basal-l and LumB-l patients, respectively (**Table 2**). By inspecting the level of phenotype activations in these subgroups we can observe that Community 2 and Community 4 are associated with increased levels of pro-oncogenic and pro-invasive phenotypes (**Figure 5C**), indicating that they might represent two subgroups of patients with the worst disease outcome. Interestingly, these two communities are enriched in Basal-l patients, suggesting that, at least, two subgroups of Basal-l patients exist and are associated with critical differences in the signaling.

Next, to demonstrate the translational relevance of our results, we used the so-obtained communities to extract relative transcriptomic signatures and benchmark them over an independent dataset. Importantly, this step aims to bridge the gap between the network-based stratifications and their potential utility in clinical settings, where transcriptomic data are widely available and easier to obtain than proteogenomic profiles.

Briefly, for each community, we considered the transcriptomic profile of the community members and, by performing an ANOVA followed by a Tukey *post-hoc* test, we identified transcripts whose up-regulation displayed the highest variance when compared to the remaining patients in the cohort. This statistical approach ensured that the selected transcripts were highly specific to the signaling rewiring characteristic of each community.

By this approach, we were able to identify seven signatures. Two of them (CL1, CL6), however, were too small and therefore were not included in the rest of the analysis.

In our cohort of Basal-l breast cancer patients, we identified two distinct groups, termed Community 2 and Community 4. To investigate the unique pathways characterizing these communities, we performed (i) an over-representation analysis of the identified signatures (**Supplementary Figure S9**), (ii) a comparison of deregulated cancer hallmarks (**Figure 5C**), and (iii) an examination of circuits driving pro-oncogenic phenotypes, at the network level (**Figure 6, and Supplementary Figure S10**). These analyses converge on two critical findings: in Community 2, there is a pronounced activation of cell growth and cell-cycle progression mediated by the MYC-CDK4/CDK6 axis, while in Community 4, a robust pro-inflammatory response is driven by the NF-kappaB pathway. At network level, we observed activation of the NF-kappaB pathway downstream of ERK1/2 and TBK1. Collectively, these findings suggest that Community 4 is associated with a metaplastic phenotype, characterized by increased aggressiveness and invasiveness^42^.

**Figure 6.**
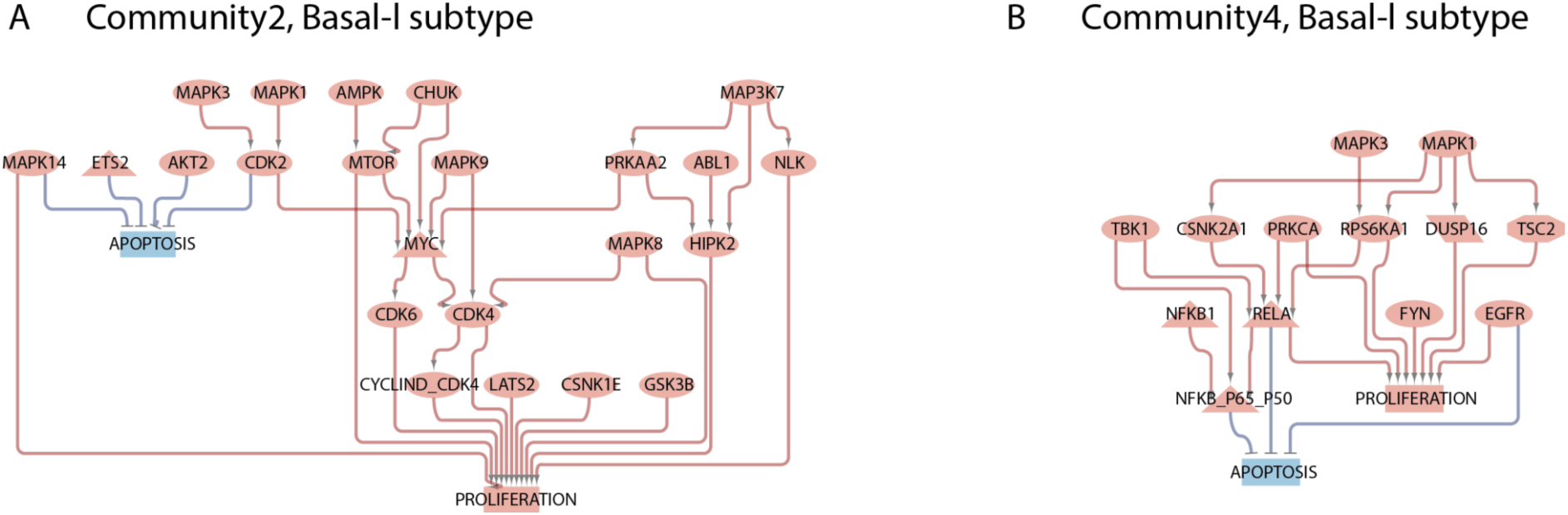
Circuits driving oncogenic phenotypes. Mechanistic models generated with *PatientProfiler*. The graph depicts the functional circuits to key phenotypes (rectangles) in A) Community 2; and B) Community 4. Red and blue nodes represent active and inactive proteins, respectively; red arrows and t-shaped edges represent activatory and inhibitory relationships, respectively.

Next, to demonstrate a broader application of our findings, we used the so-obtained signature in an independent dataset. Briefly, we accessed transcriptomic, clinical, and follow-up data from The Cancer Genome Atlas (TCGA)^41^, relative to a cohort of 1094 breast cancer patients. As described in the method section, we normalized data and, for each patient in the TCGA, we performed gene set enrichment analysis to detect the expression level of our community signatures. Next, by leveraging follow-up data we performed a survival analysis to detect whether different signatures are associated with a different expected outcome. As displayed in **Figure 5D**, the Kaplan-Meier curve indicates a strong association of Community 2 and Community 4 with a worse overall survival (OS, p-value: 0.0035), with Community 2 associated with a better prognosis. Remarkably, TCGA patients enriched by Community 2 and Community 4 are both strongly associated with Basal-l subtype classification (**Supplementary Figure S11)**.

Taken together, these results indicate that *PatientProfiler* is a tool capable of identifying novel groups of Breast cancer patients associated with differences in pathway alteration and different risks. In particular, the analysis suggests that at least two subtypes of Basal-l tumor types exist, resulting in different prognoses and easily detectable by a transcriptomic signature. The generated results clearly highlight distinct signaling axes altered in the two groups, opening novel opportunities to tailor therapeutic strategies.

## Discussion

As cancer is a signaling disease, patients with similar pathway alterations are likely to display similar evolution of the disease and clinical outcomes. Deciphering patient-specific mechanisms of cancer cell reprogramming is, therefore, pivotal in oncology to improve diagnosis and treatment. In this paper, we thoroughly present *PatientProfiler*, a computational pipeline that leverages causal interaction data to address how the genetic and molecular background of individual patients contributes to the establishment of a malignant phenotype. *PatientProfiler* is an open-source, R-based package composed of several functions that allow for multi-omic data analysis and standardization, generation of patient-specific mechanistic models of signal transduction, and extraction of network-based prognostic biomarkers.

Here, we show that *PatientProfiler* is a modular pipeline that allows users to analyze multi-omics and multi-factorial analysis as well as to choose among pre-processed proteogenomic data from more than one thousand patient-derived samples, organized in ten different tumor types, as derived from CPTAC data. In the first module, the tool enables the estimation of protein activity as well as the generation of personalized data-driven models. Briefly, the result of this first part is a collection of coherent networks portraying the molecular mechanisms underlying patient-specific disease development. Importantly, the integration of visual features and biologically relevant phenotypes simplifies interpretation and reduces noise. Remarkably, we demonstrated that phenotypes are biologically relevant and coherently reflect disease severity.

*PatientProfiler* is one of the few tools capable of delivering context-specific networks. First attempts in this direction included approaches such as pCHIPS^43^ and COSMOS+, which combine genomic, transcriptomic, and phosphoproteomic data to create contextualized networks. Unlike pCHIPS, which generates undirected graphs, *PatientProfiler* provides directed and interpretable networks able to capture the propagation of signaling events. Additionally, while COSMOS+ has been tested only in cell lines, *PatientProfiler* demonstrates clinical relevance by identifying prognostic signatures in patient-derived samples. In addition, in the case of COSMOS+ and pCHIPS, the endpoint of the pipeline is represented by the sole networks. *PatientProfiler* goes further: the second module of *PatientProfiler* allows users to leverage the collection of generated networks to identify signaling-driven communities of patients and to propose transcriptomic signatures with translational and prognostic value. We here claim that these signatures might find application in clinical settings where transcriptomic profiling of diseased samples constitutes a goal within reach^44^.

The application of *PatientProfiler* to proteogenomic profiles of Breast cancer patients, available at CPTAC, allowed us to generate a collection of 122 mechanistic networks. This collection represents, *per se*, a goldmine of information that can be used to further explore the mechanisms of tumorigenesis and cancer progression. Indeed, the individual models can be accessed for analysis, visualization, and reuse in NDEx. This collection, by displaying protein nodes according to their activity status and according to their hierarchical role in impacting cancer hallmark phenotype, might support the prioritization of personalized therapies. Furthermore, we could exploit the topological features of the so-generated graphs to detect seven communities of Breast cancer patients sharing common deregulation of signaling pathways and defined by a transcriptomic signature. These communities revealed that there might exist at least two subgroups of Basal-l, associated with distinct prognostic values.

To date, various methodologies were developed to stratify patients beyond the traditional subtypes, by generating transcriptomic signatures. These include unsupervised clustering of transcriptomic data, often coupled with dimensionality reduction techniques like PCA or t-SNE^45^; clustering methods such as iCluster, iClusterPlus, and iClusterBayes^46^; and experimental techniques such as single-cell RNA sequencing (scRNAseq) which supports the identification of cell-specific transcriptomic signatures, revealing heterogeneity within tumors and identifying rare cell populations driving disease progression^47^. Albeit powerful from the prognostic perspective, these methods are not able to provide mechanistic insight and are complementary to *PatientProfiler* in offering additional layers of biological insight and potential clinical applications.

In summary PatientProfiler is the sole tool that can generate simultaneously patient-specific directed networks with clinical relevance that can be used to priritize therapeutic targets and to generate transcriptomic signatures. These features position it as a first-of-its-kind framework that advances precision oncology beyond transcriptomic clustering.

PatientProfiler’s potential use beyond cancer (e.g., immune diseases, neurological disorders) to increase appeal.

We extensively described how *PatientProfiler* can be useful to help researchers and clinicians to extract actionable insights from their data. Despite that, it is important to stress that the tool presents limitations. The first constraint is represented by the prior knowledge space. Indeed, *PatientProfiler* suffers from the restricted coverage of available regulon and interaction information in public resources. As a matter of fact, presently *PatientProfiler* has the potential to include approximately 7,500 protein nodes (<50% of the UniprotKB proteome) and 43,145 potential regulatory interactions in a network. This is a strong underestimation with respect to the estimated number of molecular interactions occurring in cells^48^. To alleviate this burden, we implemented regular updates to ensure prior knowledge data is maintained up-to-date.

Another important limitation to mention is that, so far, *PatientProfiler* only considers transcriptomics and (phospho)proteomics, thereby narrowing down the regulatory mechanisms described in the models. Additional-omic levels such as epigenetics, metabolomics, and ubiquitylomics (among others) are becoming more common and popular^49^ and should be considered for future developments of the tool.

Both these aspects critically impact the capability of the tool to detect effective communities and, consequently, transcriptomic signatures. As shown, some signatures did not enrich patients in independent datasets.

To conclude, *PatientProfiler* addresses the emergent need to extract interpretable networks and derive biologically relevant information from complex multi-omics data and meets the long-standing challenge of generating “one model for one patient”, posing the basis for future development in the personalized medicine field. Most *PatientProfiler* is a flexible and open-source package and its applicability is not limited to cancer study.

In addition, *PatientProfiler* enables clinicians and researchers to stratify patients according to key signaling events and demonstrates that patients with similar pathway alterations are associated with different prognoses.

## Acknowledgments

This research was funded by the Italian Association for Cancer Research (AIRC) with a grant to L.P. (MFAG Grant n. 28858) and a grant to F.S. (Start-Up Grant n. 21815). Also, L.P. and F.S. are supported by a joint PRIN 2022 PNRR grant (n. P2022JRETW), funded by the European Union—NextGenerationEU, and by a SEED Sapienza Grant. V.L. is supported by a PNRR fellowship, D.M. 118/2023, funded by MUR. V.V. is supported by PON-MUR fellowship (n. DOT13IEP1U-1). M.L.N is supported by MUR - PRIN_2022 (CUP:E53D23009910006). L.D.R. is supported by Sapienza University of Rome (Grant No. RG123188B4885C78) and by INdAM – GNCS Project 2024, CUP_E53C23001670001.

## Contribution

Conceptualization, V.L., L.D.R.; L.P., F.S.; methodology, V.L., L.D.R., L.P.; formal analysis, V.L; investigation, V.L, L.P.; package development V.L., E.M., V.V., L.D.R., M.L.N.; writing original draft preparation; V.L.,V.V., L.D.R., F.S., L.P.; resources; experimental validation, E.D.N., V.P., R.N.; writing review and editing, F.S., L.P. C.C., F.S., R.N.; supervision, F.S., L.P.; funding acquisition, F.S., L.P. All authors have read and agreed to the published version of the manuscript.

## Data availability

No new experimental data was generated as part of this study. The Breast Cancer datasets are taken from ref^14^.

Networks generated are available as an NDEx collection at:

Patient-specific networks: https://www.ndexbio.org/#/networkset/bb975f9c-dcac-11ef-8e41-005056ae3c32?accesskey=5210ba80885b9ca10526b2fa52bfcf6643bb50e88401219c574796207db01b69

Communities network:

https://www.ndexbio.org/#/networkset/b03ab6d1-dcaf-11ef-8e41-005056ae3c32?accesskey=d974e5ac1f418cca58857dcf28044b9f9122c2c0e058980872e47c34f8cc380f

## Code availability

PatientProfiler R package code is available at https://github.com/SaccoPerfettoLab/PatientProfiler. The code used for the Breast Cancer use case is available at https://github.com/SaccoPerfettoLab/PatientProfiler_BRCA.

