## Supplementary Material for "*PatientProfiler:* A network-based approach to personalized medicine"

Supplementary File

Extended methods

*Step 4: Network-based patient Stratification.* This step leverages the collection of patient-specific mechanistic networks $\mathcal{G} = \{G_i, i = 1, \ldots, N\}$ to simultaneously uncover patient clusters and extract the specific pathways highly associated with them. To this end, we design a pipeline consisting of four modules: 1) Networks Aggregation. In this module, we merge the collection of networks $\mathcal{G}$ into a single aggregated network $G^*$ that stores all the interactions observed in at least one patient within the input cohort. We focus exclusively on the biological and mechanistic relevance, by excluding pathways from $G^*$originating from an inhibited node. This approach allows us to retain only functional drivers. Removing this subset of pathways ensures the analysis focuses on functionally active relationships likely to be involved in pathological regulatory mechanisms with an actual effect on the phenotype. In addition, this refinement filters out the potential background noise hindering the identification of patient communities; 2) Bipartite Projection. The second module builds a bipartite graph $B = (V, E)$ to model the patient-interaction relationship. In this graph, the vertex set is defined as $V = \mathcal{P} \cup \mathcal{I}$ , where $\mathcal{P}$ represents the set of patients in the cohort and $\mathcal{I}$ denotes the set of interactions occurring in $G^*$. An edge $e = (p, i)$ exists between a patient node $p \in \mathcal{P}$ and an interaction node $i \in \mathcal{I}$ if and only if the corresponding patient $p$ includes the interaction $i$ in their interactome. Consequently, the degree $d(i)$ of a node $i \in \mathcal{I}$ represents the number of patients whose regulatory network contains that specific interaction. This topological structure enables us to link patients with common regulatory interactions. 3) *Community Detection.* The third module involves detecting communities in $B$. To achieve this, we leverage the concept of modularity[1] a quality metric that evaluates the strength of a network's partition into clusters. Modularity is widely used for community detection, including in bipartite networks[2]. By maximizing modularity in $B$, our pipeline identifies clusters of patient nodes that are highly interconnected through subsets of shared interactions. To compute the optimal partition, we use the Louvain algorithm[3], which applies greedy optimization techniques to iteratively refine clusters. This hierarchical approach begins by grouping nodes into communities that maximize modularity. These communities are then aggregated into super-nodes, and the process is repeated on the resulting graph. The algorithm continues this iterative optimization until no further improvements or refinements can be achieved. To reduce noise during the clustering phase, our pipeline applies a preliminary filter to interaction nodes in $\mathcal{I}$, removing those with degrees below a lower threshold $t_L$ or above a upper threshold $t_U$. Indeed, interactions with low degrees could represent rare events or simply noise, lacking significance for patient stratification. Conversely, interactions with high degrees tend to be less informative, as they are common across most of the dataset. These ubiquitous interactions do not support distinguishing between patients and can hide the specific signals of the communities. For the lower threshold $t_L$, we have set a value of 4. 4) *Community Identification.* In the last module, we identify the specific pathways associated with each patient community. This is achieved by assigning to each community the set of interactions of $G^*$ grouped within the same cluster on the bipartite graph $B$ in the previous step.Indeed, the output of the third step consists of clusters of patients and the corresponding sets of interactions that make same-cluster individuals highly interconnected. These relationships can exist as isolated interactions or aggregate into larger structures, forming pathways specific to the molecular mechanisms within a  patient community.

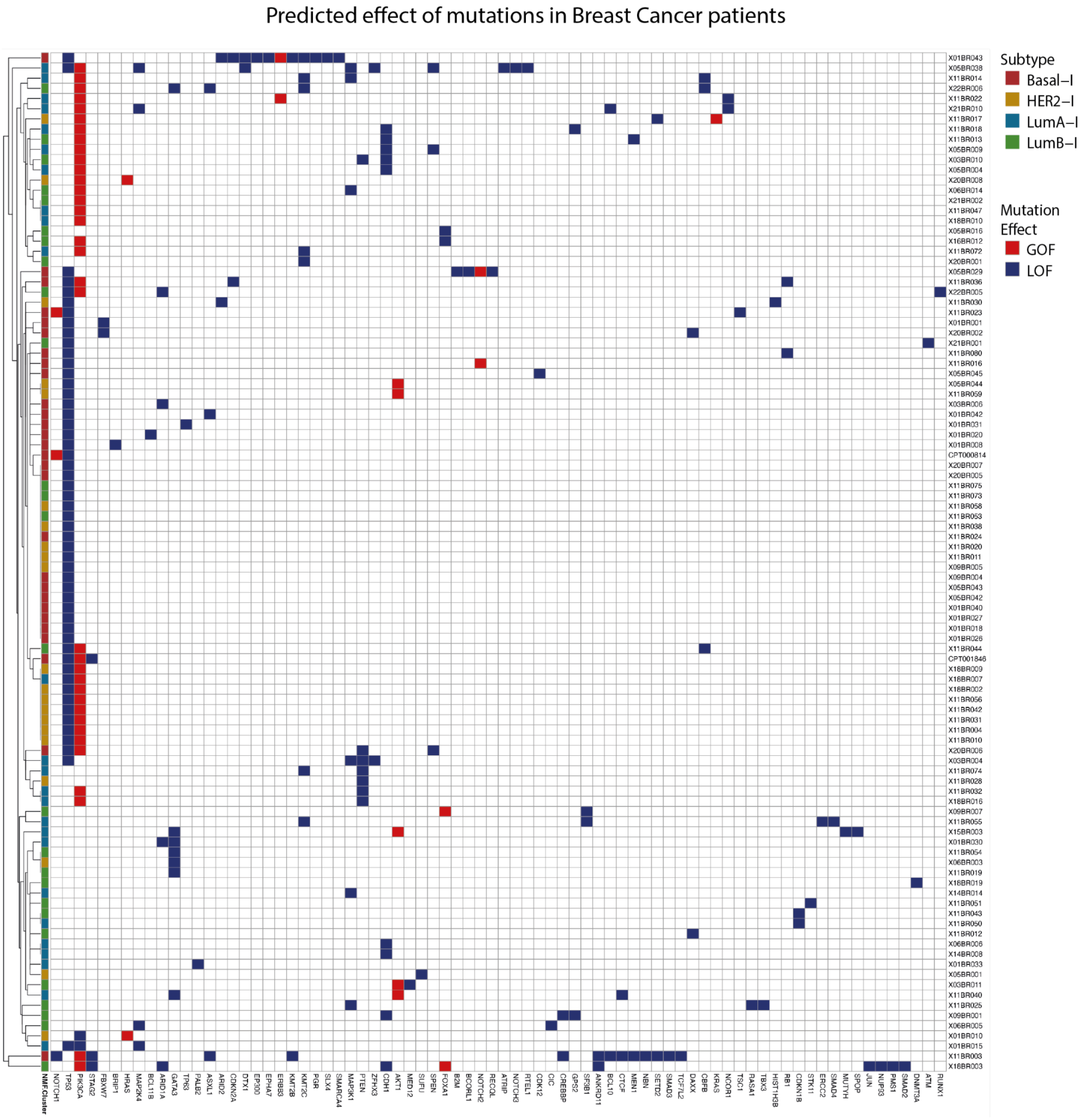


**Supplementary Figure 1 Clustering based on the mutational profile of breast Cancer Patients.** Breast cancer genomic data was retrieved from the CPTAC portal. Briefly, we exploited OncoKB (https://www.oncokb.org/)[4] to estimate the functional impact of mutations and to derive their relative impact on protein activity: loss-of-function (LOF - in blue) and gain-of-function (GOF - in red) mutations were associated to inactivation or activation of the target protein, respectively.

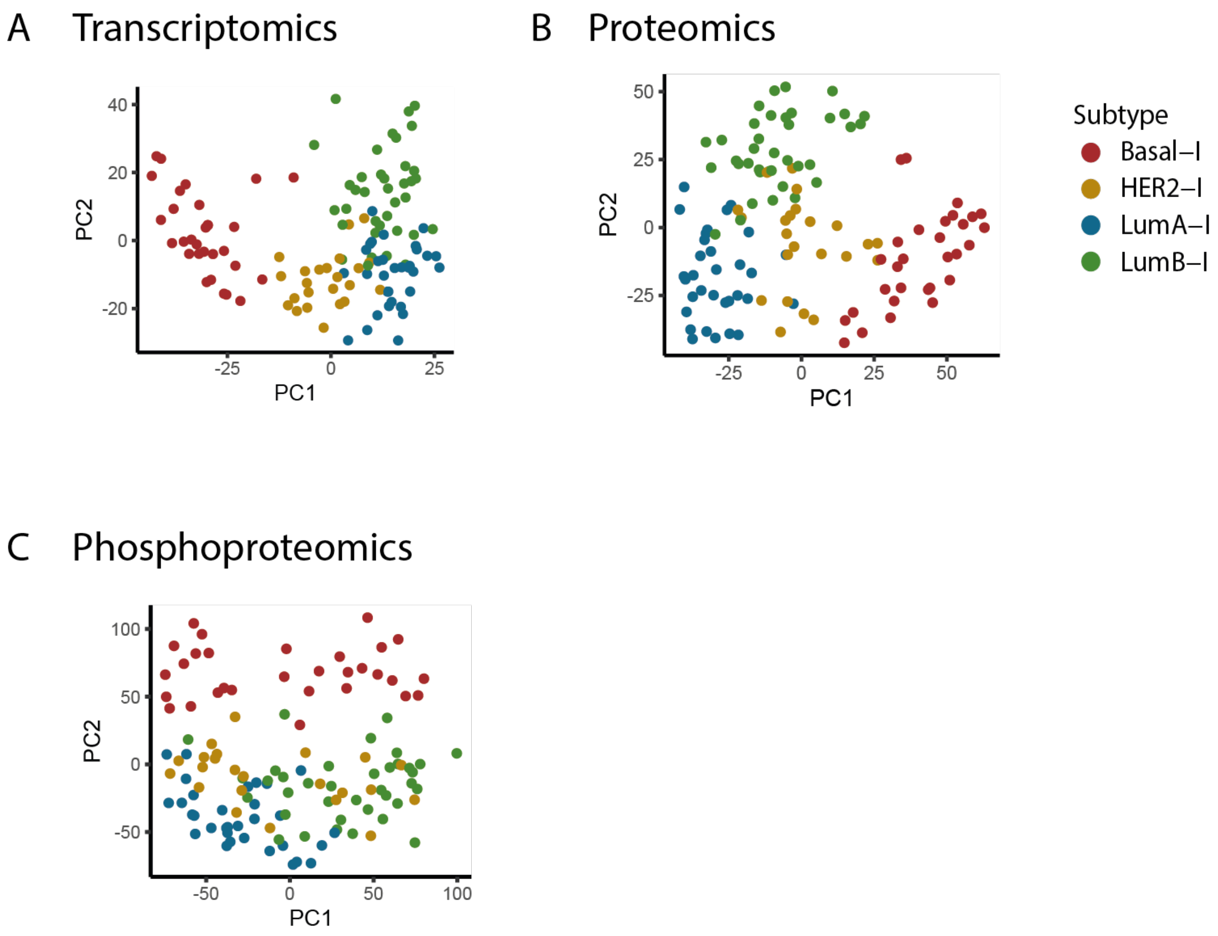


**Supplementary Figure 2. Subtype stratification at multi-omic levels.** Principal component Analysis Protein (PCA) at A) Transcriptomic; B) Proteomic; and C) Phosphoproteomic levels, after data manipulation (STEP1). Subtype color code: Basal-I in red, LumA-I in blue, LumB-I in green and HER2-I in yellow.

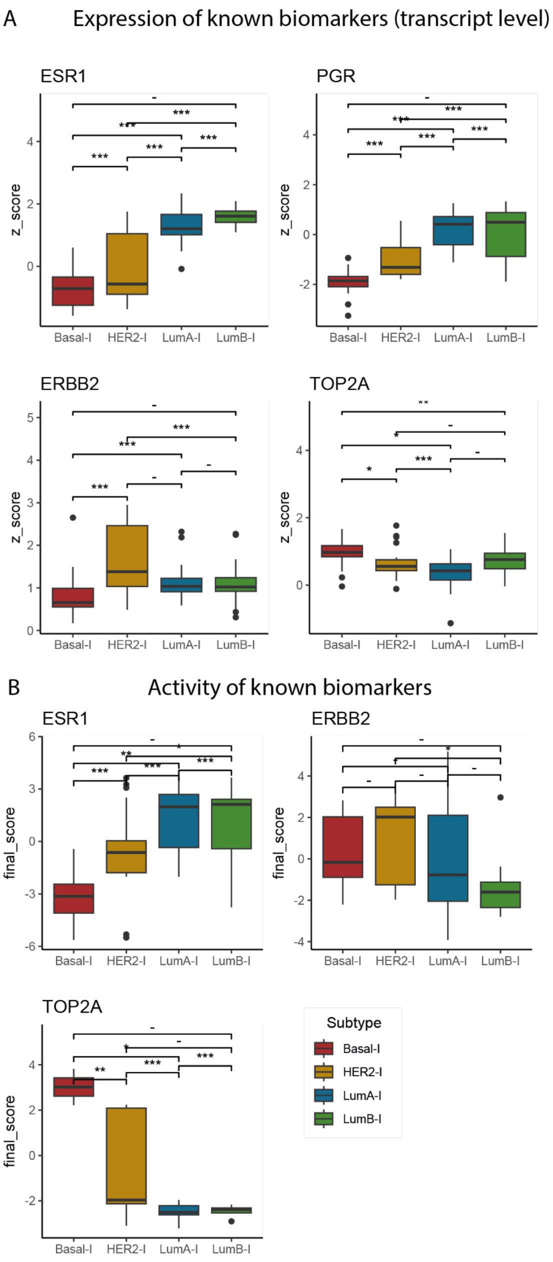


**Supplementary Figure 3 Subtypes known biomarkers. A)** Transcript expression level of known biomarkers in different subtypes, after data manipulation (STEP1). **B)** Protein activity level of known biomarkers in different subtypes, as inferred in STEP2 (ESR1: estrogen receptor, PGR: progesterone receptor, ERBB2/HER2: human epidermal growth factor receptor 2, TOP2A: proliferative biomarker). Subtype color code: Basal-I in red, LumA-I in blue, LumB-I in green and HER2-I in yellow.

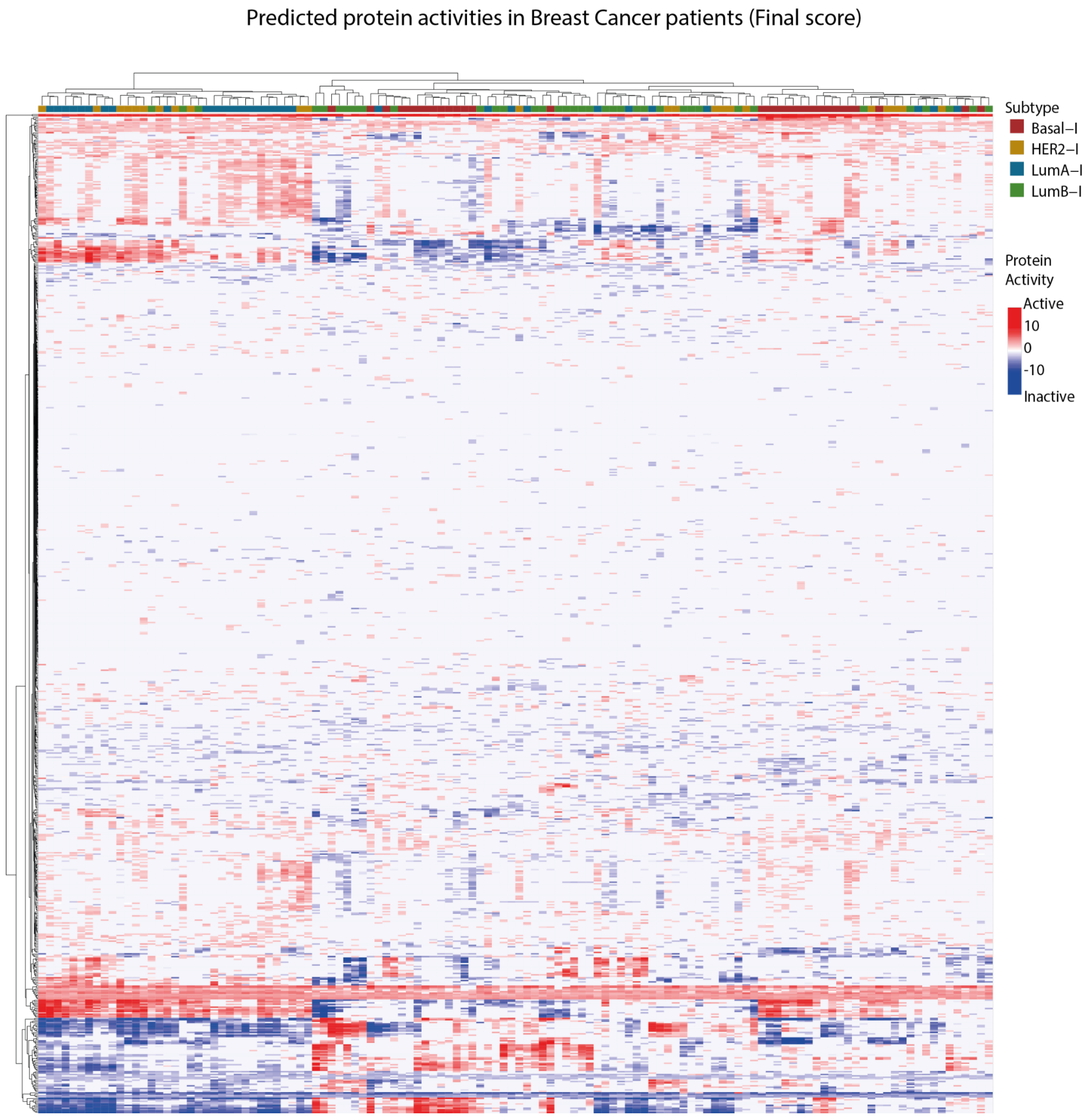


**Supplementary Figure 4 Predicted protein activities in the Breast Cancer cohort (Final score).** Heatmap showing the activation level of individual proteins in individual patients, as derived from the final score, which estimates activity from the multi-omic profiles (STEP2 of the *PatientProfiler* pipeline). Red and blue indicate active and inactive proteins, respectively. Patients and genes are clusterized (as shown in the dendrograms) accordingly, using Euclidean distance. Patients are annotated with subtype classification: Basal-I in red, LumA-I in blue, LumB-I in green and HER2-I in yellow.

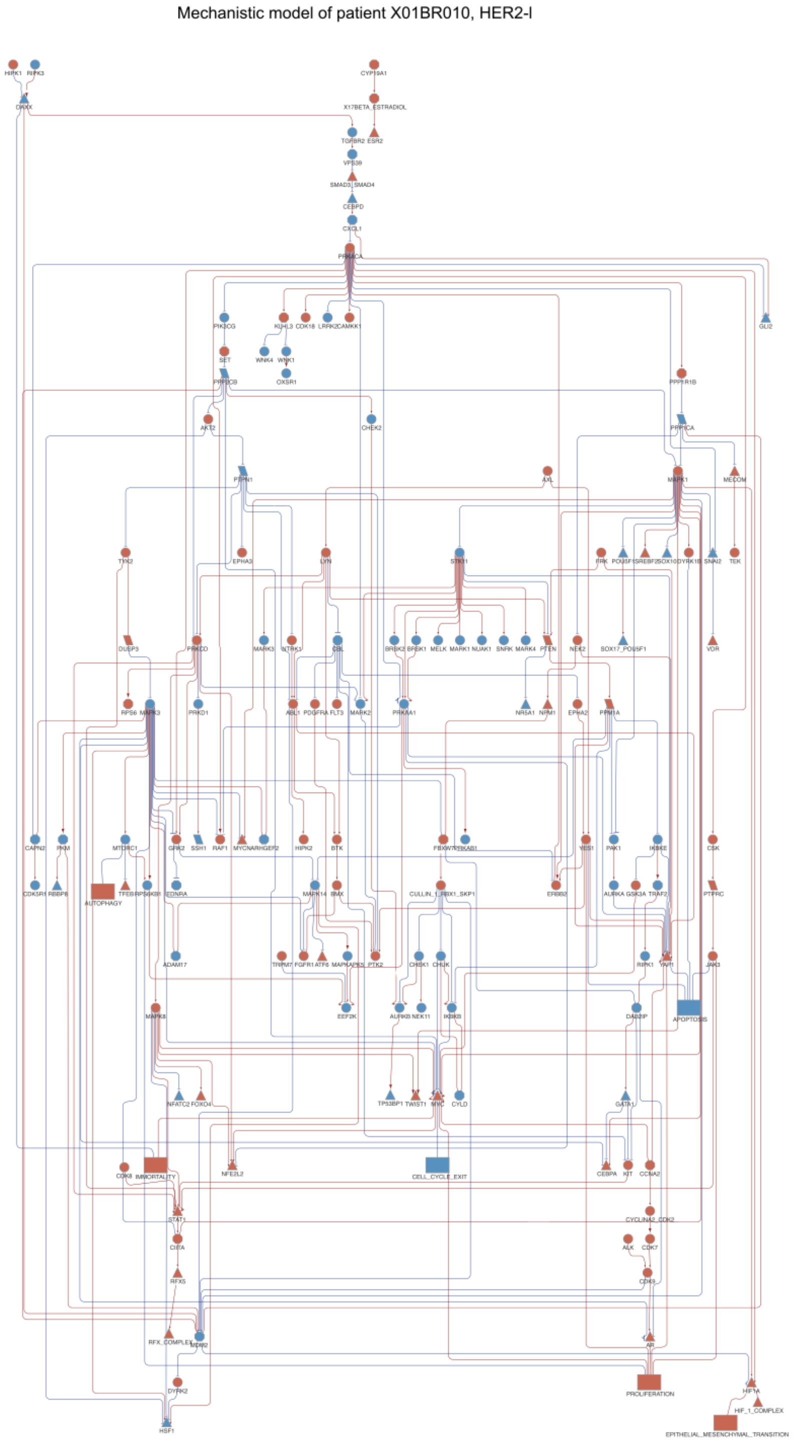


**Supplementary Figure 5 Example of a patient-specific mechanistic model.** Mechanistic model of the HER2-I patient X01BR010. Red and blue nodes represent active and inactive proteins, respectively; red arrows and t-shaped edges represent activatory and inhibitory relationships, respectively. Detected phosphorylated residues are shown as edge labels (e.g., Ser668). NDEX LINK: [https://www.ndexbio.org/#/networkset/bb975f9c-dcac-11ef-8e41-005056ae3c32?accesskey=5210ba80885b9ca10526b2fa52bfcf6643bb50e88401219c574796207db01b69](https://www.ndexbio.org/#/networkset/bb975f9c-dcac-11ef-8e41-005056ae3c32?accesskey=5210ba80885b9ca10526b2fa52bfcf6643bb50e88401219c574796207db01b69)

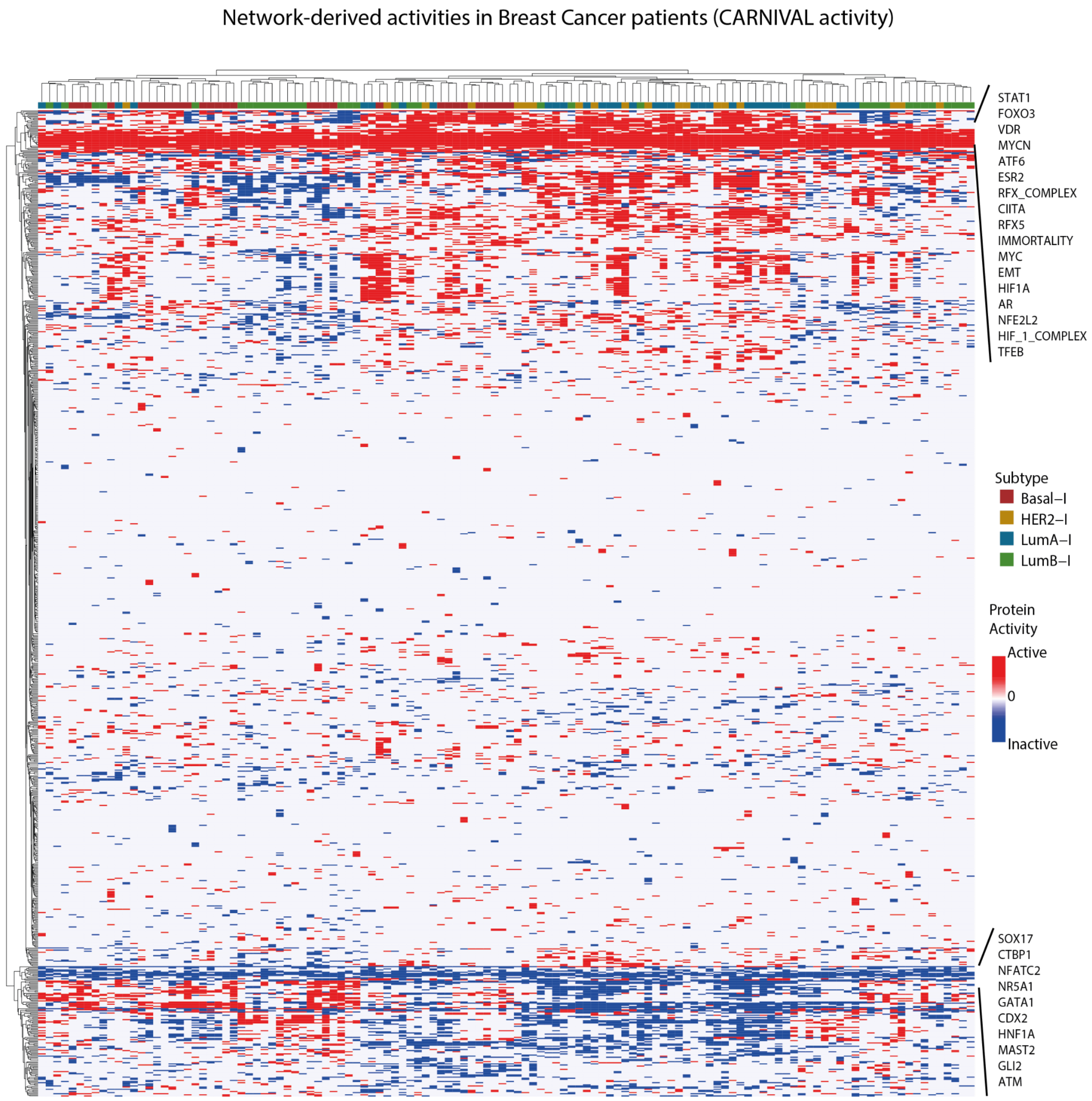


**Supplementary Figure 6 Predicted protein activities in the Breast Cancer cohort (CARNIVAL activity).** Heatmap showing the activation level of individual proteins in individual patients, as derived from the CARNIVAL score (STEP3 of the *PatientProfiler* pipeline). Red and blue indicate active and inactive proteins, respectively. Patients and genes are clusterized (as shown in the dendrograms) accordingly, using the Euclidean distance. Patients are annotated with subtype classification: Basal-I in red, LumA-I in blue, LumB-I in green and HER2-I in yellow.

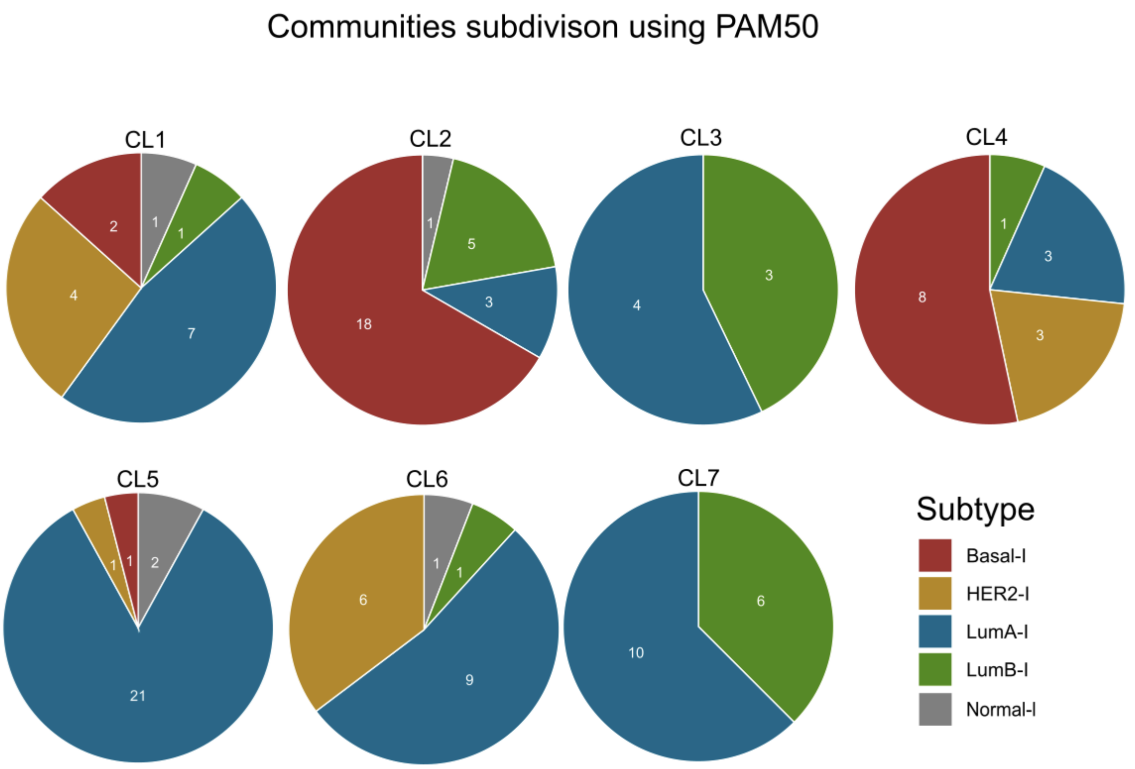


**Supplementary Figure 7 Communities' subtype composition (PAM50).** The cohort of 122 Breast Cancer patients is divided in seven communities (CL1-7). Pie charts depicting the subtype composition of each community. Subtype color code refers to the PAM50 classification[5], as derived from the original publication[6]: Basal-I in red, LumA-I in blue, LumB-I in green, HER2-I in yellow and Normal-I in gray.

**Supplementary Figure 8 Expression level of the signatures. A-G)** Plots showing the average expression level (z-score, as derived in STEP1) of each gene in each transcriptomic signature (as derived in STEP4), relative to the patients belonging to the community (CL1-7) in respect to the rest of the cohort (Other - in gray). Plots refer to (**A**) Community 1; (**B**) Community 2; (**C**) Community 3; (**D**) Community 4; (**E**) Community 5; (**F**) Community 6; and (**G**) Community 7.

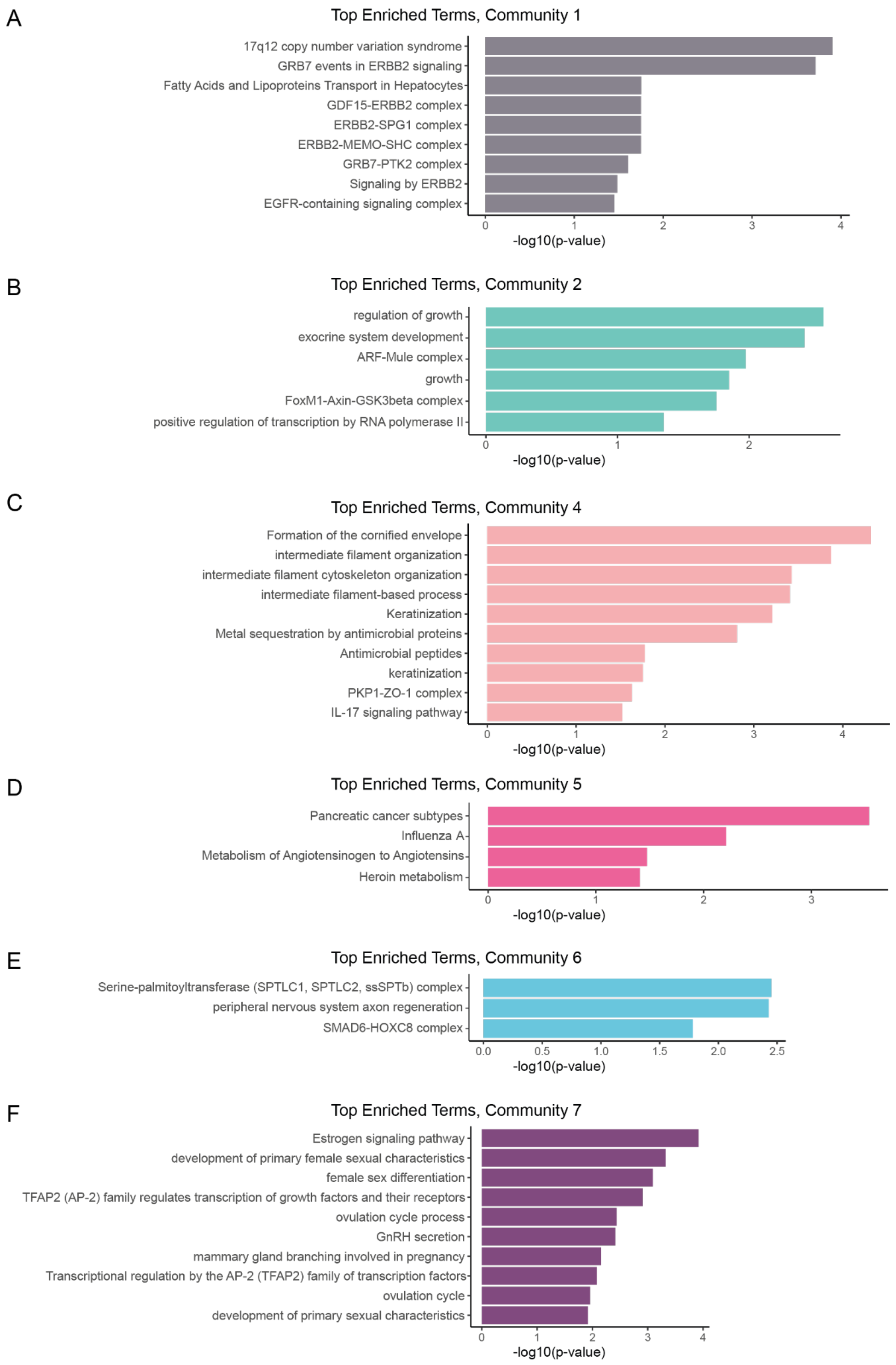


**Supplementary Figure 9 Terms overrepresentation analysis of community-derived transcriptomic signatures. A-F)** Pathway and biological process terms over-represented in each community-derived transcriptomic signature. The plots list the GO: Biological Processes, Reactome, WikiPathways and KEGG Pathways[7,8,9,10] significantly enriched in the signature. Only the top 10 significantly (Bonferroni-adjusted *P*-value < 0.05) over-represented terms are shown. Analyses were carried out by using the gProfiler2 software[11], using the entire Human Proteome as a background. Plots are relative to (**A**) Community 1; (**B**) Community 2; (**C**) Community 4; (**D**) Community 5; (**E**) Community 6; (**F**) Community 7.

A Community2, 191 nodes and 242 edges

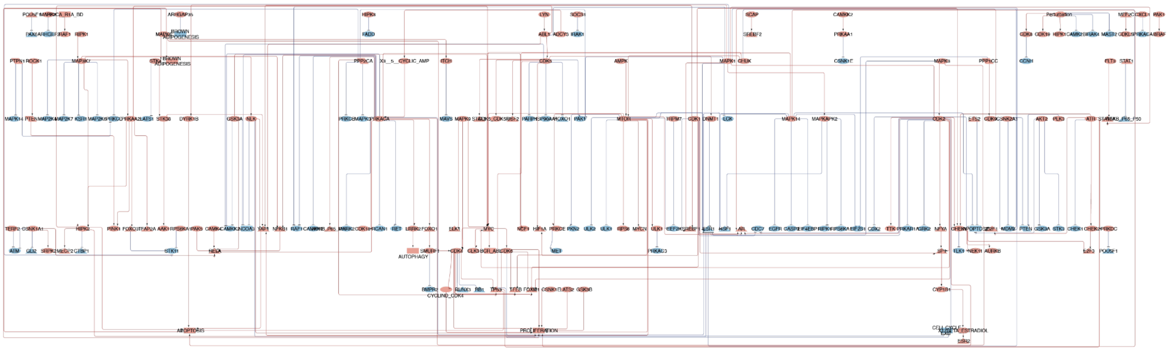

B Community4, 120 nodes and 131 edges

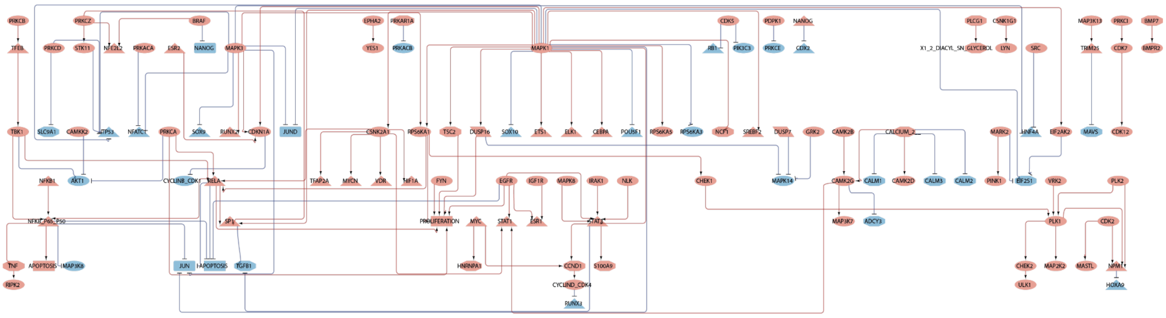

**Supplementary Figure 10 Basal-I communities .** Mechanistic models generated with *PatientProfiler*. The graph depicts the functional graph associated with in **A)** Community 2; and **B)** Community 4. Red and blue nodes represent active and inactive proteins, respectively; red arrows and t-shaped edges represent activatory and inhibitory relationships, respectively. NDEX LINK: [https://www.ndexbio.org/#/networkset/b03ab6d1-dcaf-11ef-8e41-005056ae3c32?accesskey=d974e5ac1f418cca58857dcf28044b9f9122c2c0e058980872e47c34f8cc380f](https://www.ndexbio.org/#/networkset/b03ab6d1-dcaf-11ef-8e41-005056ae3c32?accesskey=d974e5ac1f418cca58857dcf28044b9f9122c2c0e058980872e47c34f8cc380f)

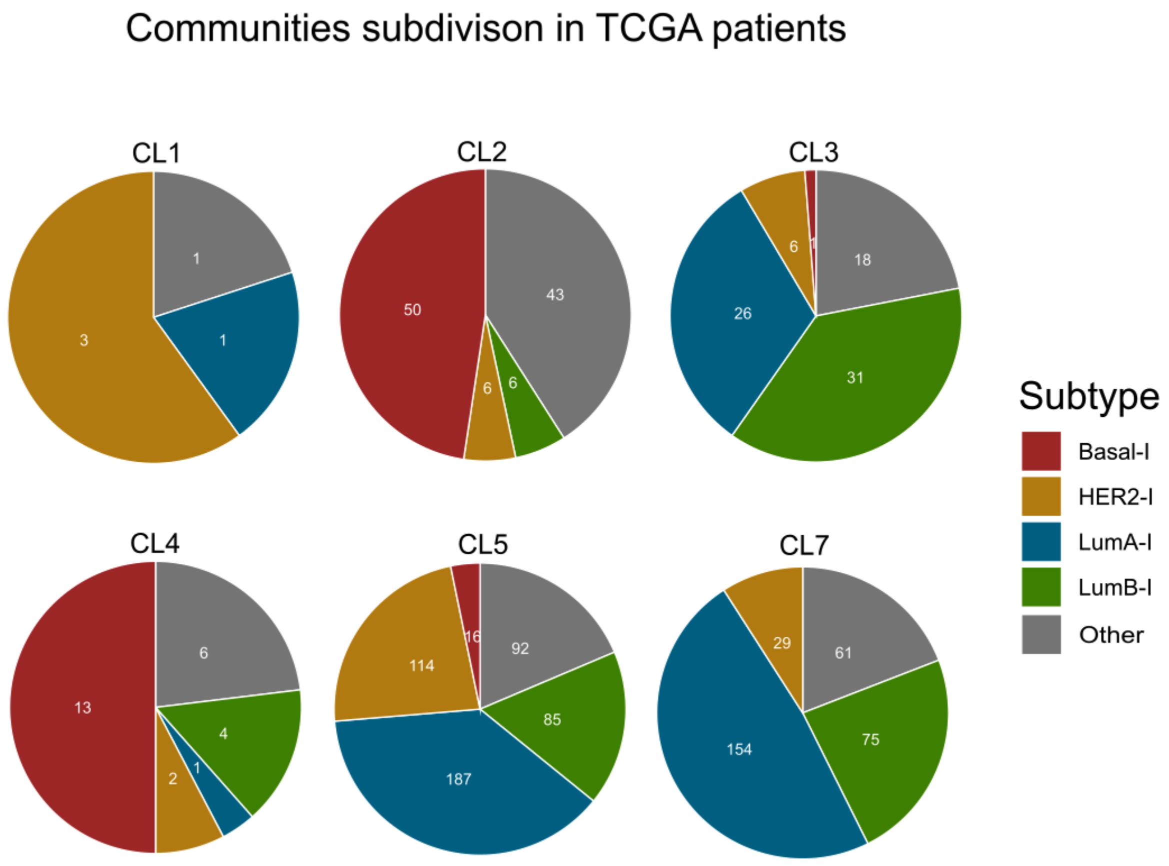


**Supplementary Figure 11 Community-based stratification of TCGA Breast Cancer patients.** Breast Cancer patients-derived transcriptomic data from The Cancer Genome Atlas (TCGA)[12] was used in combination (gene set enrichment analysis - GSEA) with community-derived transcriptomic signatures to stratify TCGA patients in seven communities (CL1-7) (Adjusted P-value < 0.01, NES > 0 for CL2, CL3, CL4, CL7, NES > 1 for CL1, CL5, CL6). Pie charts depicting the subtype composition of each community. Subtype color code: Basal-l in red, LumA-l in blue, LumB-l in green, HER2-l in yellow and Other (unclassified) in Gray.
